# S-Nitrosylation of CRTC1 in Alzheimer’s disease impairs CREB-dependent gene expression induced by neuronal activity

**DOI:** 10.1101/2023.08.25.554320

**Authors:** Xu Zhang, Roman Vlkolinsky, Chongyang Wu, Nima Dolatabadi, Henry Scott, Olga Prikhodko, Mayra Blanco, Nhi Lang, Juan Piña-Crespo, Tomohiro Nakamura, Marisa Roberto, Stuart A. Lipton

## Abstract

CREB-regulated transcription coactivator 1 (CRTC1) plays an important role in synaptic plasticity, learning and long-term memory formation through regulation of neuronal activity-dependent gene expression, and CRTC1 dysregulation is implicated in Alzheimer’s disease (AD). Here, we show that increased S-nitrosylation of CRTC1 (forming SNO-CRTC1), as seen in cell-based, animal-based, and human induced pluripotent stem cell (hiPSC)-derived cerebrocortical neuron-based AD models, disrupts its binding with CREB and diminishes the activity-dependent gene expression mediated by the CRTC1/CREB pathway. We identified Cys216 of CRTC1 as the primary target of S-nitrosylation by nitric oxide (NO)-related species. Using CRISPR/Cas9 techniques, we mutated Cys216 to Ala in hiPSC-derived cerebrocortical neurons bearing one allele of the APP^Swe^ mutation (AD-hiPSC neurons). Introduction of this non-nitrosylatable CRTC1 construct rescued defects in AD-hiPSC neurons, including decreased neurite length and increased neuronal cell death. Additionally, expression of non-nitrosylatable CRTC1 *in vivo* in the hippocampus rescued synaptic plasticity in the form of long-term potentiation (LTP) in 5XFAD mice. Taken together, these results demonstrate that formation of SNO-CRTC1 contributes to the pathogenesis of AD by attenuating the neuronal activity-dependent CREB transcriptional pathway, and suggests a novel therapeutic target for AD.

## INTRODUCTION

Protein S-nitros(yl)ation occurs when nitric oxide (NO)-related species, possibly as transition-metal catalyzed transfer of a nitrosonium cation (NO^+^) intermediate, react with the thiolate anion (RS^-^) of a target cysteine residue to form a SNO-protein^1^. This reaction has emerged as an important posttranslational modification (PTM) reminiscent of *O*-phosphorylation^2–4^. Physiological protein S-nitrosylation in the brain has been implicated in regulation of cellular processes such as ion channel activity, synaptic plasticity, neuronal development, and cell survival^5,6^. However, under neurodegenerative conditions such as Alzheimer’s disease (AD), excessive generation of reactive nitrogen species (RNS), including NO-related molecules, are triggered by amyloid-β peptide oligomers (Aβo), neuronal hyperexcitability, or neuroinflammation. This results in aberrant S-nitrosylation of many proteins that would not normally manifest the SNO PTM^7–9^. Such aberrant protein S-nitrosylation can impact enzyme activity, protein localization, and protein–protein interactions, thus contributing to the pathogenesis of neurodegenerative disorders^7–9^.

Activity-dependent gene expression is important for neuronal plasticity and memory consolidation. Disruption of gene-expression has been postulated to occur at early stages of AD and altered gene expression in the brain is thought to precede the neuropathological changes of AD^10–13^. The transcription factor, cAMP response element-binding protein (CREB), has been shown to be essential for long-term synaptic plasticity, neurite outgrowth, memory consolidation and neuronal survival^14–17^. Dysregulation of CREB-dependent gene expression has been implicated in early stages of AD, contributing to synaptic dysfunction^18–20^. Additionally, CREB regulated transcription coactivator 1 (CRTC1) plays a critical role in regulating activity-dependent gene expression as a coactivator of CREB^21,22^. There are three isoforms of CRTCs in mammalian cells, and CRTC1 is the major isoform in neurons^23^. Unlike CREB, which is constitutively located in the nucleus, CRTC1translocates into the nucleus in response to electrical activity^24^, and thus represents a form of signal transduction from synapses and dendrites to the nucleus.

Under basal conditions, CRTC1 is phosphorylated by salt-inducible kinases1 and 2 (SIK1/2) and binds to 14-3-3 protein, which sequestrates CRTC1 in the cytosol^24,25^. Synaptic activation promotes Ca^2+^ influx and increases cAMP levels, which in turn activates calcineurin and protein kinase A (PKA), respectively. PKA phosphorylates and thus inhibits SIK1/2 kinase, thus inhibiting CRTC1 phosphorylation. Additionally, calcineurin dephosphorylates CRTC1, resulting in CRTC1 dissociating from 14-3-3 and translocating into the nucleus^24,26^. Once in the nucleus, CRTC1 binds in a complex to CREB and chromatin, thus contributing to expression of CREB targets, such as the immediate-early genes (IEGs). Among these IEGs, *cfos*, *erg1* (early growth response protein 1), and *Arc* (activity-regulated cytoskeleton-associated protein) are thought to represent master regulators of synaptic plasticity^24,27^. Accordingly, CRTC1 contributes to dendritic outgrowth of developing cerebrocortical neurons, and knockout of CRTC1 decreases expression of several genes involved in neuroplasticity^28,29^. Studies of postmortem human brain tissue have shown that CRTC1 protein levels and associated gene transcription are significantly reduced in the human hippocampus of AD patients with intermediate-to-advanced Braak stages (IV-VI) AD^30^. Moreover, transgenic mouse models of AD that overexpress human amyloid precursor protein (hAPP) exhibit impaired expression of CRTC1-dependent genes at early stages of disease prior to CRTC1 depletion^10,11,30^, suggesting there are other, yet unknown, molecular mechanisms that regulate CRTC1 activity.

Here, we show that excessive levels of NO, as seen in AD and other neurodegenerative disorders, can S-nitrosylate CRTC1 to disrupt its interaction with CREB, thus inhibiting activity-dependent CRTC1/CREB-mediated gene expression. We found that SNO-CRTC1 is elevated under neurodegenerative conditions in cell-based models and in brain lysates of transgenic AD mice, and we identify Cys216 as the primary target of S-nitrosylation. We then show that S-nitrosylation of CRTC1 following Aβo-induced generation of NO disrupts the interaction of CRTC1 with CREB in rat primary cerebrocortical cultures. This results in decreased expression of CRTC1/CREB regulated genes. We use CRISPR/Cas9 to knock-in a Cys216 to Ala mutation in cell-based models, including human induced pluripotent stem cell (hiPSC)-derived cerebrocortical neurons with one allele bearing the APPSwe mutation (AD-hiPSC neurons). We observed that SNO-CRTC1 contributes to dysregulated gene expression and morphological defects in these AD-hiPSC neurons compared to controls, while the non-nitrosylatable CRTC1^C216A/C216A^ mutant can largely reverse these defects. Finally, we utilized the 5XFAD transgenic AD mouse model to investigate the effects of lentiviral expression of non-nitrosylatable CRTC1^C216A^ vs. wild-type (WT) CRTC1 in the hippocampus. Our results demonstrate that overexpression of CRTC1^C216A^, but not WT CRTC1, significantly increased synaptic markers in 5XFAD mice. Additionally, impaired long-term potentiation (LTP) in 5XFAD mice was rescued by overexpression of CRTC1^C216A^ in the hippocampus. These findings have important implications for disease-modifying therapy in AD.

## RESULTS

### S-Nitrosylation of CRTC1 at Cys216

Initially, we tested if endogenous CRTC1 can be S-nitrosylated (forming SNO-CRTC1) by exposing rat primary cerebrocortical cultures to the physiological NO donor and transnitrosylating agent S-nitrosocysteine (SNOC). As a control, cells were exposed to ‘old’ SNOC from which NO had been dissipated. Ten minutes later, to detect SNO-CRTC1 we performed a biotin-switch assay as previously described^31,32^. In this assay protein free thiols are blocked with methyl methanethiosulfonate (MMTS) and S-nitrosothiols are specifically reduced by ascorbate to generate free thiols. The newly generated thiols are labeled with biotin and pulled down with NeutrAvidin beads. Subsequently, immunoblotting is performed to analyze the pulled-down samples. Using this method, we observed a significant increase in SNO-CRTC1 levels in primary cerebrocortical neurons after incubation with SNOC **(Figure 1a).** We also overexpressed all three isoforms of CRTC in HEK293 cells and used biotin-switch assays to detect the various SNO-CRTC isoforms after exposure to SNOC. We found that all three CRTC isoforms can be S-nitrosylated (**Extended Data Figure 1a**). CRTCI, however, is the predominant isoform in the brain, and particularly in the hippocampus^23^. Moreover, CRTC1 affects CREB activity, which is critical for memory formation and is altered in AD^21,22,30^. Hence, we decided to study in detail the effects of S-nitrosylated CRTC1.

**Figure 1.**
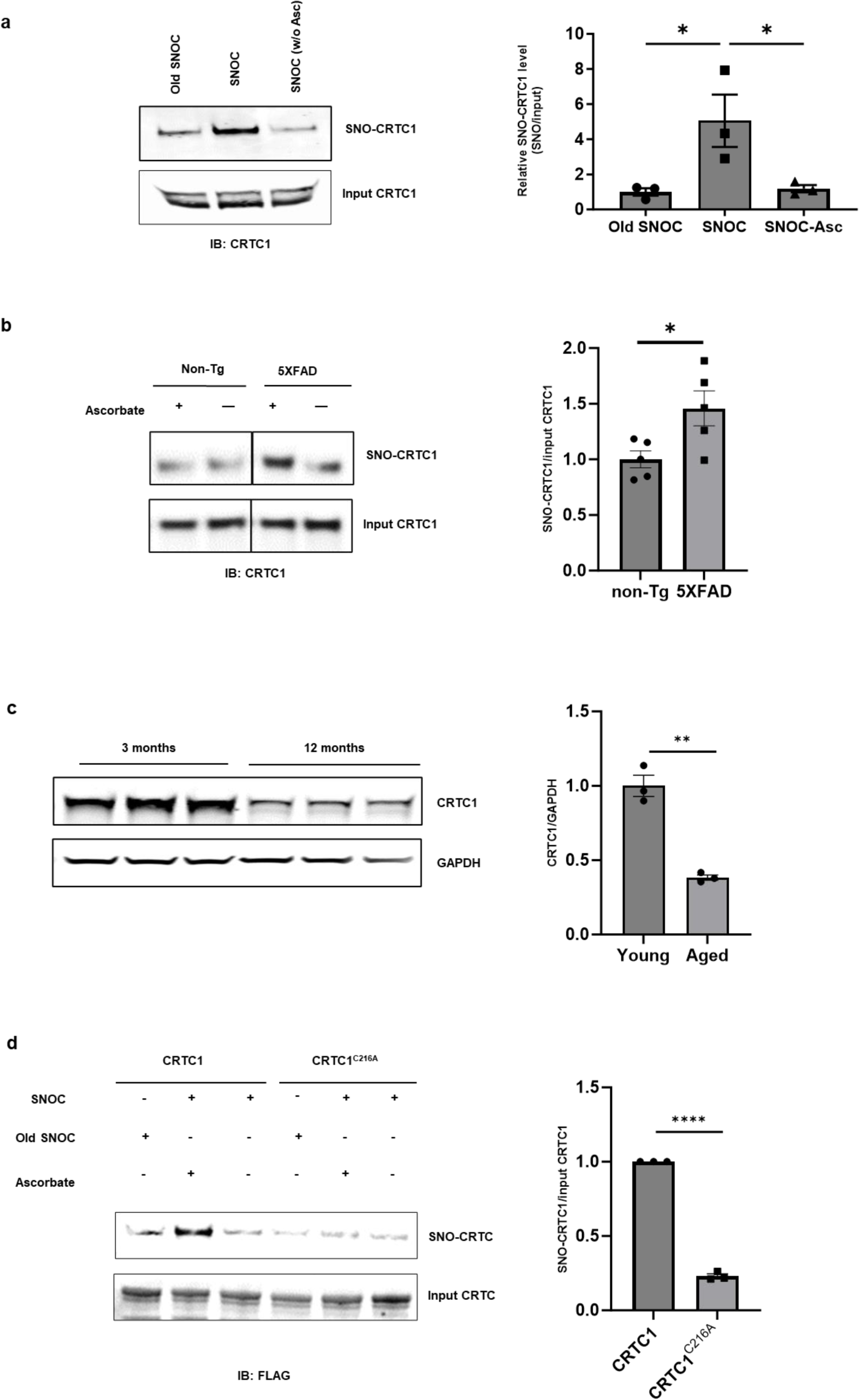
S-Nitrosylation of endogenous CRTC1 in primary rat cerebrocortical cultures and transgenic AD mice. **a,** Formation of SNO-CRTC1 in cerebrocortical neurons. Neurons were exposed to SNOC (100 µM) or, as a control, equimolar “old SNOC” (from which NO had been dissipated). S-Nitrosylation of endogenous CRTC1 (SNO-CRTC1) was assessed by biotin-switch assay. SNOC-exposed samples in the absence of ascorbate (w/o Asc) served as a negative control. Note that multiple bands of CRTC isoforms have been reportedly previously on immunoblotting in the loading controls^82,83^, as observed here in the CRTC1 loading controls. Values are mean ± SEM; n = 3 independent experiments, analyzed by ANOVA followed by Tukey’s multiple comparisons test, **p < 0.01. **b,** Presence of SNO-CRTC1 in the cerebrocortex of 5XFAD mouse brain. Brain lysates from 3∼4-month-old 5XFAD or non-Tg (WT) littermate mice were subjected to biotin-switch assay. Ratio of SNO-CRTC1/input-CRTC1 expressed as mean ± SEM; n = 5 mice per genotype analyzed by two-tailed Student’s t test, *p < 0.05. **c,** Decreased CRTC1 levels in aged mouse brain. Brain lysates from 3-month-old (young) and 12-month-old (aged) WT mice were separated by SDS-PAGE and analyzed by immunoblot. Ratio of CRTC1/GAPDH expressed as mean ± SEM; n = 3 mice per age group analyzed by two-tailed Student’s t test, **p < 0.01. **d,** Cys216 is the predominant site of S-nitrosylation in CRTC1. SH-SY5Y cells were transfected with FLAG-fused constructs encoding WT or C216A mutant CRTC1 and subjected to biotin-switch assay. Ratio of SNO-CRTC1/input-CRTC1 expressed as mean ± SEM; n = 3 independent experiments analyzed by two-tailed Student’s t test, ****p < 0.0001.

To test if SNO-CRTC1 is elevated under neurodegenerative conditions *in vivo*, we analyzed brain lysates from the 5XFAD transgenic (Tg) AD mouse model, which expresses five mutations in human APP and PS1 (5XFAD) (APP^SweFlLon^, PSEN1^*M146L*L286V^). Age-matched WT littermates were used as controls (non-Tg). By 3∼4 months of age, biotin-switch assays revealed a significant increase in SNO-CRTC1 in 5XFAD mouse brain lysates compared to non-Tg controls (**Figure 1b**). At this time point, the disease is just beginning to be manifest by behavioral, electrophysiological, and histological criteria^33^. Similarly, we also observed elevated level of SNO-CRTC1 in 3 months old hAPP-J20 mouse brain lysates compared to age-matched non-Tg controls (**Extended Data Figure 1b**). When we tried to confirm the presence of increased levels of SNO-CRTC1 in human AD brain tissues, however, the protein level of CRTC1 in both AD patient and age-matched control frontal lobe tissue was barely detectable in postmortem samples; this of course could have reflected the state of the human postmortem tissue (**Extended Data Figure 1c**). In agreement with this finding, however, aged mice also manifest greatly decreased CRTC1 levels by 12 months of age (**Figure 1c**). Taken together, these observations are consistent with the notion that the detection of SNO-CRTC1 may represent an early biomarker in the AD brain, which is not evident at later stages of the disease (e.g., when postmortem human brain tissue becomes available) because of very low levels of CRTC1 protein in general in aged brain. Additionally, dysregulation of CRTC1-dependent transcription in the brains of the hAPP-J20 mouse and other transgenic models has been reported starting at early stages of AD-related damage^10,30^. These data motivated us to study if S-nitrosylation contributes to aberrant CRTC1 activity early in the development of the neurodegenerative disease process.

Initially, we determined the cysteine residue(s) involved in SNO-CRTC1 formation using site-directed mutagenesis in HEK-293 cells. CRTC1 contains only 2 free candidate cysteine thiols^22^. Additionally, there is only one homologous cysteine residue conserved across all three mammalian CRTCs: Cys216 in CRTC1, Cys245 in CRTC2 and Cys244 in CRTC3 (**Extended Data Figure 1d**). Indeed, we found that mutation at CRTC1^C216A^ decreased S-nitrosylation by ∼80% compared to WT CRTC1 (**Figure 1d**). We also mutated Cys245 in CRTC2 and found that it is the predominant site of S-nitrosylation in CRTC2 as well (**Extended Data Figure 1e**).

### S-Nitrosylation triggers CRTC1 nuclear translocation in a Ca^2+^- and calcineurin-dependent manner

Next, we studied the effect of the NO donor/transnitrosylating agent SNOC on activity-dependent translocation of CRTC1 from spines and dendrites into the nucleus of excitatory neurons. This translocation is known to require the calcium- and calmodulin-dependent serine/threonine protein phosphatase, calcineurin^24^, which dephosphorylates CRTC1. Since increased NO is known to disrupt Ca^2+^ homeostasis in neurons^34,35^, we investigated if the subcellular distribution of CRTC1 is altered in primary rat cerebrocortical neurons in response to SNOC. We exposed rat cortical neurons to 100 µM SNOC, and after 5 min analyzed the cellular distribution of CRTC1 by immunohistochemistry. We found that SNOC promoted nuclear translocation of CRTC1. This translocation was blocked by the intracellular Ca^2+^ chelator BAPTA-AM); inhibition of calcineurin with cyclosporin A also prevented CRTC1 transport into the nucleus (**Figure 2a**). These results are consistent with the notion that Ca^2+^ and calcineurin are key players in NO-triggered nuclear translocation of CRTC1.

**Figure 2.**
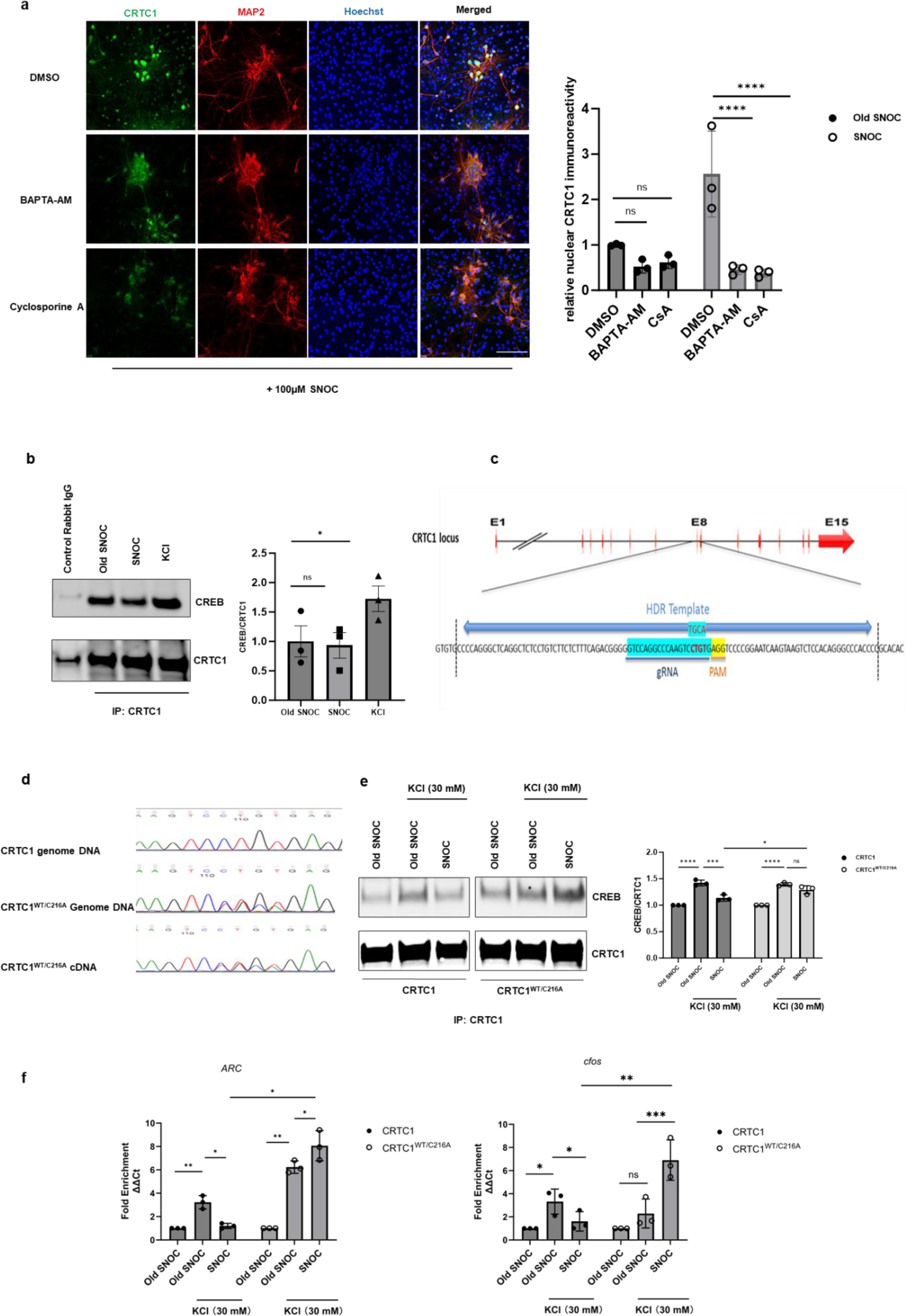
S-Nitrosylation of CRTC1 disrupts its interaction with CREB. **a,** Cellular distribution of CRTC1 after SNOC exposure. The membrane-permeable calcium chelator BAPTA-AM (25 µM, 30 min) and the calcineurin antagonist cyclosporine A (10 µM, 1 h) were incubated with primary rat cerebrocortical cultures prior to SNOC exposure. Neurons were then fixed and immunostained with primary antibodies against MAP2 (red), CRTC1 (green), and Hoechst nuclear dye (blue). Quantitative CRTC1 nuclear immunoreactivity was analyzed by two-way ANOVA followed by Tukey’s multiple comparisons test, n = 3 samples, ****p < 0.001, ns: not significant. Scale bar, 100 µm. **b,** Co-immunoprecipitation of CRTC1 and CREB in primary rat cerebrocortical cultures 5 min after exposure to SNOC (100 μM). Incubation in KCl (30 mM, 5 min) was used as a positive control to induce CRTC1 binding to CREB. Co-immunoprecipitations were performed with anti-CRTC1 antibody or, as a control, rabbit isotype-specific IgG. Western blots were probed with anti-CRTC1 and anti-CREB antibodies. Ratio of CREB/CRTC1 binding expressed as mean ± SEM; n = 3 independent experiments analyzed by ANOVA, and Tukey’s multiple comparisons test, *p < 0.05. **c,** Design of the HDR template, target location and target sequences of the CRISPR/Cas9 system used in SH-SY5Y cells. **d,** Target mutation of C216A in the CRTC1 gene was confirmed by Sanger sequencing. **e,** Co-immunoprecipitation of CRTC1 and CREB shows differential abilities of CRTC1^WT/WT^ and mutant CRTC1^WT/C216A^ to bind to CREB. SH-SY5Y cells were pre-exposed to SNOC (100 µM). After 5 min, cells were washed, allowed to recover for 1 hour, and then stimulated with KCl (30 mM, 5 min). Cells were then collected and subjected to co-immunoprecipitation with anti-CRTC1 antibody. Ratio of CREB/CRTC1 binding expressed as mean ± SEM; n =3 independent experiments analyzed by two-way ANOVA and Tukey’s multiple comparison test, **p < 0.01, ***p < 0.001, ****p < 0.0001, and ns: not significant. **f,** Quantitative ChIP-qPCR analysis shows differential recruitment of CRTC1^WT/WT^ and mutant CRTC1^WT/C216A^ to the promoter regions of c*fos* and *Arc* after SNOC and KCl incubation. SH-SY5Y cells were treated in the same way as in panel (E), then cells were collected and subjected to ChIP-qPCR. Data represent mean fold enrichment (−ΔΔCT) ± SEM; n = 3 independent cultures from 3 independent experiments analyzed by two-way ANOVA followed by Tukey’s multiple comparisons test, **p < 0.01, ***p < 0.001, ****p < 0.0001, and ns: not significant.

### Nuclear translocation of CRTC1 is not affected by S-nitrosylation of CRTC1 per se

Next, we investigated whether S-nitrosylation of CRTC1 affects its nuclear translocation. For this purpose, we exposed primary rat cerebrocortical cultures to 100 µM SNOC or control solution. Five minutes later, after washing the cells, we placed them back into the conditioned medium (which had been collected prior to SNOC exposure) and allowed the cells to recover for one hour. During this recovery period, CRTC1 was transported out of the nucleus, as monitored by immunostaining (**Extended Data Figure 2a**), but S-nitrosylation of CRTC1 was maintained, as verified by biotin-switch assay (**Extended Data Figure 2b**). In contrast, treatment with 30 mM KCl 1-hr post SNOC exposure triggered transport of CRTC1 back into the nucleus (**Extended data Figure 2a**).

### S-Nitrosylation of CRTC1 disrupts the interaction of CRTC1 with CREB

Following nuclear translocation, CRTC1 is known to interact with CREB to facilitate transcription of downstream genes; thus, CRTC1 acts as a transcriptional co-regulator. To test the possible effect of S-nitrosylation of CRTC1 on this process, we immunoprecipitated CRTC1 from rat cerebrocortical neurons to detect CRTC1 binding to CREB in the presence and absence of SNOC. After exposure to SNOC, CRTC1 accumulated in the nucleus (**Figure 2a),** but there was no increase CRTC1/CREB binding (**Figure 2b**). As a positive control, we produced activity-dependent translocation of CRTC1 and an increase in CRTC1/CREB binding by depolarizing the cells with 30 mM KCl for 5 minutes (**Figures 2b**). As a further control, treatment with 30 mM KCl alone for 5 minutes did not result in the formation of SNO-CRTC1 (**Extended Data Figure 2c**). Taken together, these results are consistent with the notion that formation of SNO-CRTC1 disrupts the binding of CRTC1 to CREB, while Ca^2+^ accumulation via depolarization by KCl or intracellular mobilization via NO, triggers calcineurin-dependent dephosphorylation to effect transfer of CRTC1 into the nucleus.

To test the effect of SNO-CRTC1 on the interaction of CRTC1 and CREB further, we used CRISPR/Cas9-mediated gene editing to introduce a single amino acid mutation (C216A) into the CRTC1 gene, initially using the human neural cell line SH-SY5Y. By nucleofection, we transduced the cells with a plasmid carrying the Cas9 gene and guide (g)RNA targeting the editing site together with a single-stranded oligodeoxynucleotide (ssODN) homologous directed repair (HDR) template (**Figure 2c**). We then isolated single cells by fluorescence-activated cell sorting (FACS), followed by the generation of single-cell clones. Sanger sequencing was used to confirm the identity of the knock-in mutation and verified that we had generated heterozygous colonies bearing WT and C216A mutant (CRTC1^WT/C216A^) alleles (**Figure 2d**). We also obtained single-cell colonies without mutation of the target site (CRTC1^WT/WT^) and used these as controls. Heterozygous mutation did not detectably change the expression level of CRTC1, since the WT and mutant alleles are both functional (**Extended Data Figure 3a**). As expected, however, we observed a dramatic decrease in SNO-CRTC1 formation (**Extended Data Figure 3b**).

Next, we performed co-immunoprecipitation experiments to determine if the activity-dependent interaction of CRTC1 with CREB was affected by SNO-CRTC1. For this experiment, we exposed CRTC1^WT/WT^ and CRTC1^WT/C216A^ cells to 100 µM SNOC or control solution; after 5 minutes, we washed the cells and placed them back into conditioned medium collected prior to SNOC (or control) (exposure in order to allow them to recover for 1 h. This recovery period allowed CRTC1 to be transported back out of the nucleus (as demonstrated in **Extended Data Figure 2a**), while maintaining its S-nitrosylation (**Extended Data Figure 2b**). We then incubated the cells for 5 min in high-K^+^ depolarizing solution. Depolarization with high K^+^ produced nuclear translocation of WT CRTC1 in cells previously exposed to SNOC or control “old” SNOC prior to collection for co-immunoprecipitation assay. This experiment confirmed that formation of SNO-CRTC1 disrupted activity (high K^+^)-dependent interaction of CRTC1 with CREB in CRTC1^WT/WT^ cells (**Figure 2e**). In contrast, cells bearing non-nitrosylatable mutant CRTC1 displayed significantly more activity-dependent binding of CRTC1 to CREB after exposure to SNOC (**Figure 2e**). As a control, activity-dependent binding of CRTC1 to CREB in the absence of SNOC was not significantly different between CRTC1^WT/WT^ and CRTC1^WT/C216A^ (**Figure 2e**), consistent with the notion that the non-nitrosylatable mutant did not affect the binding of CRTC1 to CREB per se.

To confirm that the activity-dependent interaction of CRTC1 with CREB and chromatin was disrupted by formation of SNO-CRTC1, we performed quantitative chromatin immunoprecipitation (ChIP)-qPCR. This set of experiments was carried out under the same conditions as the co-immunoprecipitation assays and showed that formation of SNO-CRTC1 disrupted activity-dependent recruitment of CRTC1 to the CREB-binding promoter region of genes such as c*fos* and *Arc* in CRTC1^WT/WT^ cells (**Figure 2f)**. In contrast, mutant CRTC1^WT/C216A^ cells, which manifest decreased S-nitrosylated CRTC1 (**Extended Data Figure 3b**), displayed an increase in CRTC1/CREB-mediated gene expression in response to KCl stimulation after SNOC exposure compared to CRTC1^WT/WT^ (**Figure 2f**). Intriguingly, in mutant CRTC1^WT/C216A^ compared to CRTC1^WT/WT^ cells, high KCl had a larger effect on *Arc* and *cfos* transcription, and SNOC further increased, rather than decreased expression of these genes (**Figure 2f**). This result was to be expected, however, because increased levels of NO had previously been shown to activate CREB activity under conditions favoring formation of SNO-CREB^36^. Moreover, this effect would be magnified in our case in the presence of non-nitrosylatable CRTC1 because the absence of SNO-CRTC1 could not diminish expression of these CREB-dependent genes^37^. Additionally, S-nitrosylation of other proteins might also have influenced the expression of CREB-dependent genes under these conditions. For example, formation of S-nitrosylated HDAC2 in the presence of BDNF has been shown to increase expression of CREB target genes^38^.

### Aβo trigger SNO-CRTC1 formation and decreases CRTC1/CREB-mediated gene expression in rat primary cerebrocortical neurons

To determine the effect of SNO-CRTC1 on gene expression in a disease-relevant model system, we exposed primary cerebrocortical neurons to Aβo, which have previously been shown to produce pathological levels of NO^39,40^. By biotin-switch assay, we found that incubation in 500 nM Aβo_1-42_ for 3 hours induced formation of SNO-CRTC1 (**Figure 3a**). When the NO synthase (NOS) inhibitor L-N^G^-nitroarginine methyl ester (L-NAME) was used to inhibit NO generation, SNO-CRTC1 formation decreased (**Figure 3a**). Additionally, pre-incubation in Aβo decreased the activity-mediated interaction of CRTC1 and CREB induced by exposure to high K^+^, but this effect was abrogated by pre-treatment with L-NAME (**Figure 3b**). As a control, incubation in Aβo alone in the absence of high K^+^ did not induce CRTC1 translocation into the nucleus (**Extended Data Figure 4**). These results are consistent with the notion that Aβo-induced S-nitrosylation of CRCT1 disrupted the interaction of CRTC1 and CREB only after neuronal stimulation with KCl had induced activity-dependent translocation of CRTC1 into the nucleus. Accordingly, ChIP-qPCR analysis of the CREB downstream genes c*fos* and *Arc* revealed that Aβo exposure significantly decreased neuronal activity-dependent transcription of these genes induced by high K^+^. Moreover, pre-treatment with L-NAME prevented the effect of Aβo (**Figure 3c**).

**Figure 3.**
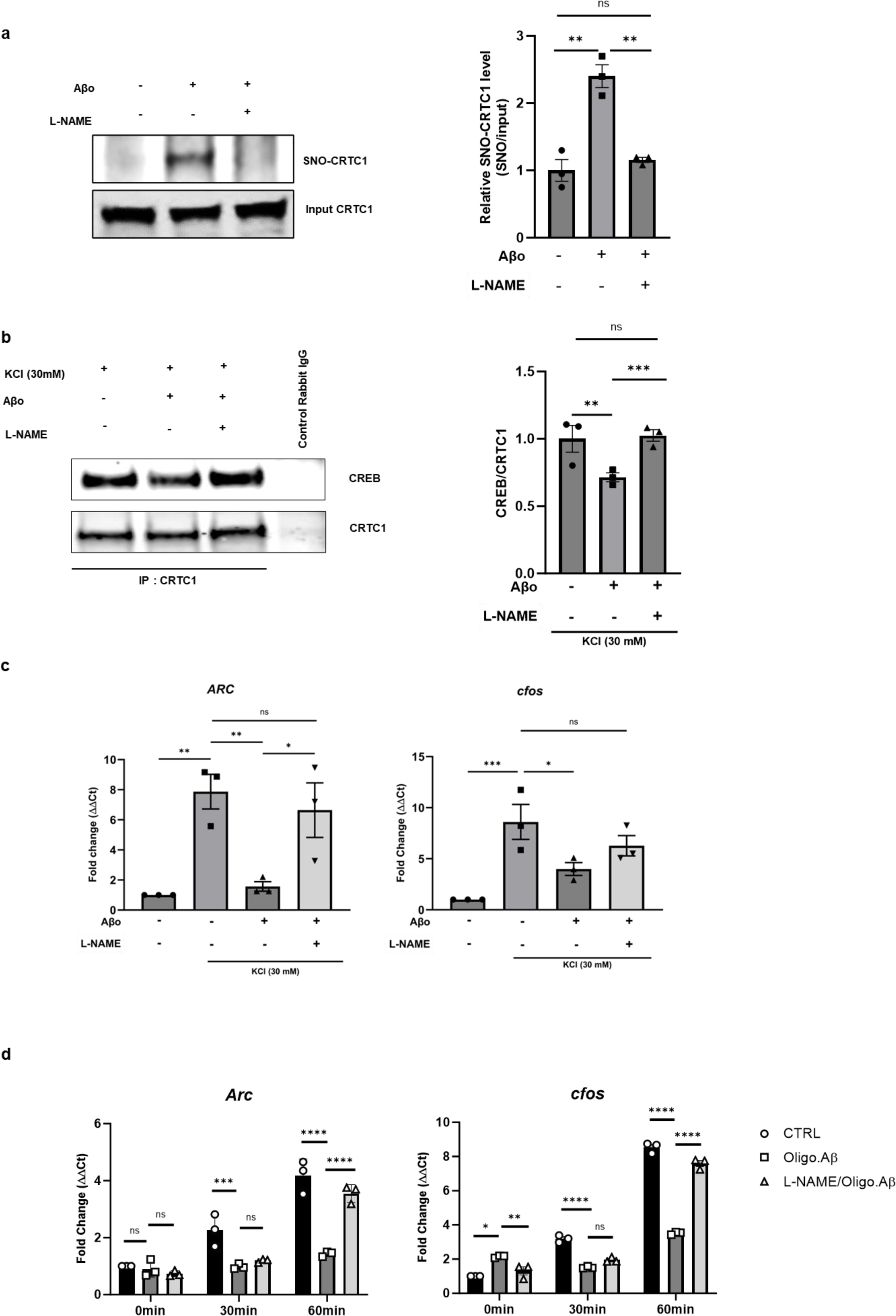
Aβo disrupt CRTC1 and CREB interaction in primary rat cerebrocortical cultures. **a,** Formation of SNO-CRTC1 in neurons exposed to Aβo (500 nM, 3 hours). NOS activity was inhibited with L-NAME (1 mM in the presence of 1 mM arginine) as a negative control. SNO-CRTC1 was detected by biotin-switch assay. Ratio of SNO-CRTC1 and input CRTC1 expressed as mean ± SEM; n = 3 independent experiments). **p < 0.01 by ANOVA and Tukey’s multiple comparisons test. **b,** Aβo decrease neuronal activity induced CRTC1/CREB binding. Neurons were incubated in 500 nM Aβo for 3 hours, and then stimulated with KCl (30 mM for 5 min). NOS activity was inhibited by pre-treatment with L-NAME (1 mM for 3 hours). Cells were then collected and subjected to co-immunoprecipitation. Ratio of CREB/CRTC1 expressed as mean ± SEM; n = 3 independent experiments analyzed by ANOVA and Tukey’s multiple comparisons test, **p < 0.01, ***p <0.001. **c,** Quantitative ChIP-qPCR analysis on KCl-depolarized neurons exposed to Aβo. Differential recruitment of CRTC1 to the promoter regions of cfos and Arc after Aβo exposure, shown as mean fold-enrichment (−ΔΔCT) ± SEM; n =3 from independent cultures analyzed by ANOVA and unpaired Fisher’s LSD test, *p < 0.05. **d,** Time-dependent increase in *cfos* and *Arc* mRNA by qRT-PCR after KCl depolarization is impaired by Aβo. Data represent mean ± SEM from n = 3 independent experiments analyzed by two-way ANOVA and Tukey’s multiple comparisons post hoc test, *p < 0.05, **p < 0.01, ***p < 0.001, and ****p <0.0001.

To further investigate the effect of SNO-CRTC1 on gene transcription activated by CREB, we performed quantitative (q)PCR to analyze the mRNA levels of c*fos* and *Arc* at two different time points: 30 and 60 minutes after KCl depolarization in the presence or absence of Aβo. We confirmed that neuronal activity (high KCl)-induced expression of the genes was significantly impaired by Aβo at both time points (**Figure 3d**). Pre-treatment with L-NAME significantly prevented the decrement in mRNA expression of both genes, but only at the 60 min time point (**Figure 3d**), probably reflecting the time-dependence of both transcriptional activation by CREB and NO generation from nNOS after Aßo exposure. Taken together, these findings are consistent with the notion that S-nitrosylation of CRTC1 leads to decreased recruitment of CRTC1 to the promoter region of CREB target genes, with resultant suppressed expression of downstream genes.

### AD-hiPSC neurons display elevated levels of SNO-CRTC1

Net, to test the effect of SNO-CRTC1 in a human context, we used AD-hiPSC neurons. We have previously shown that these neurons in culture manifest hyperexcitability, increased Aβ production, and decreased neurite length compared to isogenic, gene-corrected WT control neurons (non-AD-hiPSC neurons)^41^. NO levels have been reported to increase in human patient brains during the progression of AD, in part a result of elevated levels of nNOS and iNOS with a concomitant decline in antioxidant capacity^8,42^. Thus, we initially investigated if RNS are increased in AD-hiPSC neurons compared to non-AD-hiPSC neurons. For this purpose, we used the Griess reaction to measure total nitrite and nitrate in the conditioned medium of 4 to 6-week-old neurons. The results showed a significant increase in total nitrite/nitrate levels in AD-hiPSC neurons compared to non-AD-hiPSC neurons at the 6-week time point. Treatment with L-NAME significantly reduced the level of total nitrite/nitrate in the AD-hiPSC neurons, but not in non-AD-iPSC neurons, indicating that NO synthase was the major source of NO in the AD-hiPSC neurons. (**Figure 4a**). Accordingly, western blot analysis of cell lysates from 4-to 6-week-old cultures revealed elevated expression of nNOS in AD-hiPSC neurons compared to non-AD-hiPSC neurons (**Figure 4b**). This observation agrees with the previous finding that neurons in human AD brain aberrantly express high levels of nNOS, whereas neurons in non-pathological conditions usually express low levels of nNOS^7,43^. Next, we performed biotin-switch assays on the 6-week-old AD-hiPSC and non-AD-hiPSC neurons and found, as expected, that the ratio of SNO-CRTC1/total CRTCI was significantly increased in AD-hiPSC neurons compared to WT controls at this age (**Figure 4c**). Notably, this ratio was similar in the 5XFAD and hAPP-J20 transgenic mouse brains as in the AD-hiPSC neurons, indicating that our cell culture model manifested pathophysiologically-relevant levels of SNO-CRTC1 similar to those observed in the early stages of *in vivo* AD models (*cf.* **Figures 1c, Extended Data Figure 1b, and 4c**).

**Figure 4.**
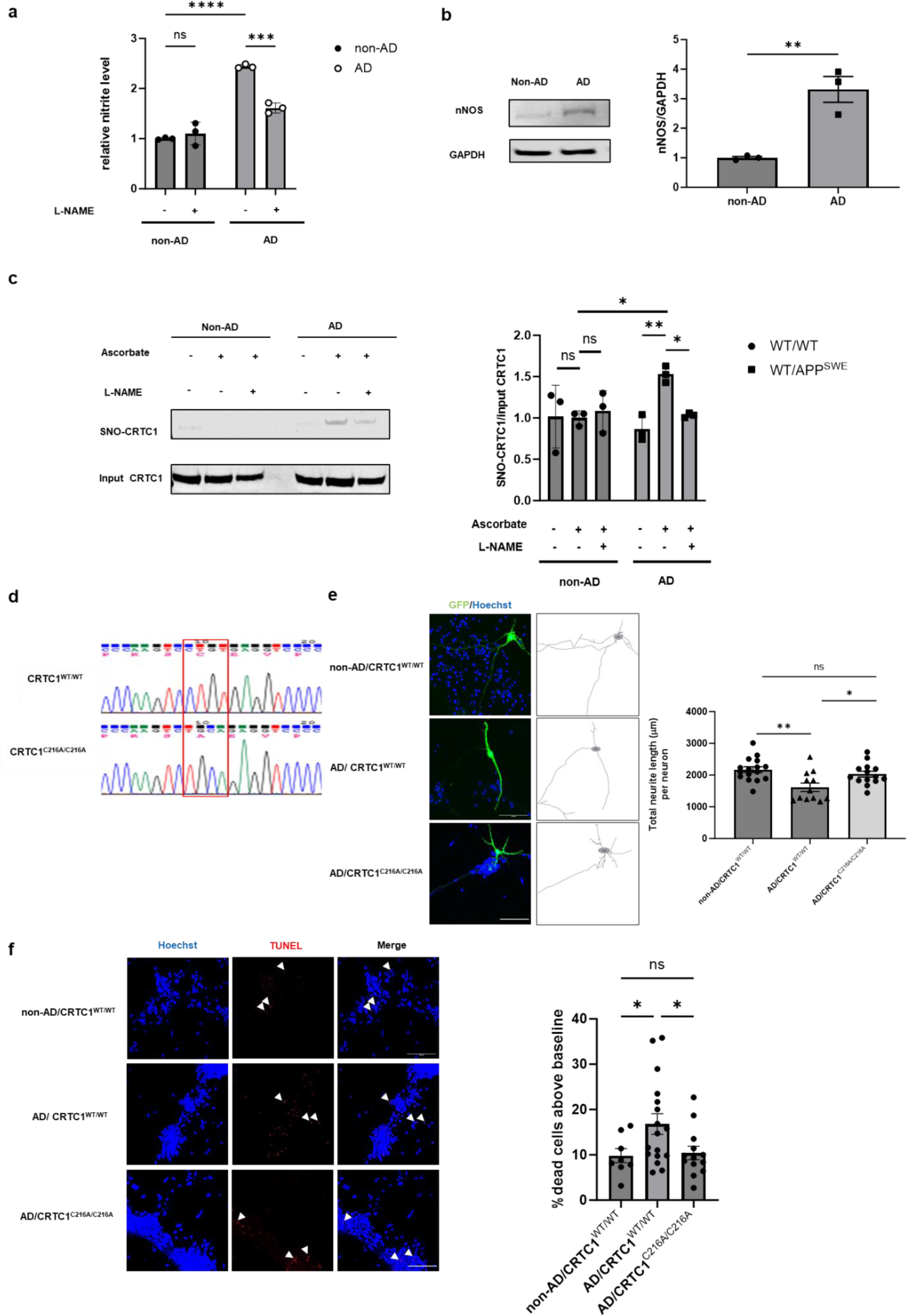
SNO-CRTC1 contributes to neuronal injury in hiPSC-derived AD neurons. **a,** Increased level of NO-related species in hiPSC-derived AD neurons. After 4-6 weeks in culture and 3 days after the last half-medium exchange, conditioned medium from hiPSC-derived AD or WT neurons was collected, and nitrite/nitrate levels determined by Griess reagent (expressed as mean ± SEM). Significant differences between AD and WT neurons were analyzed by two-way ANOVA and Turkey’s multiple comparisons post hoc test, n = 3 independent differentiations, ***p < 0.001, and ****p <0.0001. **b,** Increased expression level of nNOS in hiPSC-derived AD neurons. At the same time as in panel A, hiPSC-derived AD and WT neurons were collected, and their cell lysates subjected to immunoblot. Protein levels of nNOS were detected with anti-nNOS antibody, normalized to GAPDH, and expressed as mean ± SEM. Significance was analyzed by two-tailed Student’s t test, n = 3 independent differentiations, **p <0.01. **c,** Increased level of SNO-CRTC1 in hiPSC-derived AD neurons relative to WT neurons. Six-week-old hiPSC-derived AD and WT neuronal lysates were subject to biotin-switch assay. Difference in ratio of SNO-CRTC1/input-CRTC1 (expressed as mean ± SEM) were analyzed by two-way ANOVA and post hoc Fisher’s LSD, n = 3 independent differentiations, *p < 0.05, **p < 0.01. **d,** Sanger sequencing confirmed introduction of C216A mutation in the CRTC1 gene by CRISPR/Cas9 technology. **e,** CRTC1^C216A/C216A^ mutation abrogated neurite length defects in AD neurons. Six-week-old hiPSC-derived cortical neurons were transfected with green fluorescent protein (GFP). GFP-labeled neurons were used for measurement of total neurite length. Statistical differences between total neurite length per neuron (expressed as mean ± SEM) were analyzed by ANOVA and Tukey’s multiple comparisons post hoc test, *p < 0.05, **p < 0.01, n=3 independent differentiations, 3-7 neurons were used for neurite tracing from each differentiation (n = 16 for WT neurons, n = 12 for WT CRTC1-AD neurons, and n = 14 for mutant CRTC1^C216A/C216A^ AD neurons). Scale bar, 100 µm. **f,** CRTC1^C216A/C216A^ mutation showed decreased apoptosis compared to WT CRTC1-AD neurons by TUNEL assay. Six-week-old hiPSC-derived cortical neurons were analyzed by TUNEL staining. The percentage of TUNEL-positive cells was determined from the number of TUNEL-positive cells vs. total cell number (identified by DAPI staining) in each image. Images were obtained from 3 independent differentiations. Statistical differences between mean % TUNEL-positive cells were analyzed by ANOVA followed by Fisher’s LSD post hoc test, *p< 0.05 (n = 8 for WT neurons, n = 16 for WT CRTC1-AD neurons and n = 13 for CRTC1^C216A/C216A^ AD neurons). Scale bar, 100 µm.

### Generation of AD hiPSCs bearing non-nitrosylatable mutant CRTC1 using CRISPR/Cas9

CREB-mediated gene transcription normally fosters neurite outgrowth and neuronal survival^44^. To test the hypothesis that the reduced capacity of SNO-CRTC1 to promote CREB-mediated gene transcription affects neurite outgrowth and neuronal survival, we generated a non-nitrosylatable CRTC1 knock-in using CRISPR/Cas9. For this purpose, we electroporated AD hiPSCs with the template HDR ssODN and a ribonucleoprotein (RNP), consisting of the Cas9 protein in a complex with a gRNA targeting the knock-in site. We isolated the electroporated cells by fluorescence-assisted cell sorting (FACS) followed by single-cell clone generation and colony expansion. We then used Sanger sequencing to confirm successful editing (**Figure 4D**). Moreover, sequencing the top 3 most probable off-target sites in genomic DNA revealed no off-target editing. We characterized the newly generated hiPSC line (AD/CRTC1^C216A/C216A^) for pluripotency after generation of embryoid bodies (EBs) and performed immunohistochemistry for specific markers of germ layer generation (**Extended Data Figure 5a**). Dual-SMAD inhibition allowed us to efficiently generate neural progenitor cells (hNPCs), as confirmed by immunofluorescent labeling with the NPC markers Nestin, PAK6 and SOX2 (**Extended Data Figure 5b**). We then differentiated these AD/CRTC1^C216A/C216A^ hNPCs into cerebrocortical neurons^41^, as confirmed by staining for microtubule associated protein MAP2 and FOXG1 (**Extended Data Figure 5c**).

Next, as suggested by our prior experiments on rat primary cerebrocortical neurons, we found in the hiPSC-derived neurons that non-nitrosylatable CRTC1 had no effect on KCl-induced nuclear translocation of CRTC1. Quantification of the immunocytochemical images showed that isogenic non-AD/CRTC1^WT/WT^, AD/CRTC1^WT/WT^, and AD/CRTC1^C216A/C216A^ non-nitrosylatable mutant hiPSC-derived neurons all induced nuclear translocation of CRTC1 to a similar degree after a 5-min exposure to 30 mM KCl (**Extended Data Figure 5d**).

### Non-nitrosylatable mutant CRTC1 prevents shorter neurites and cell death in AD-hiPSC neurons

To assess differences in neuronal morphology in human cerebrocortical neurons derived from isogenic AD/CRTC1^WT/WT^ hiPSCs, non-AD/CRTC1^WT/WT^, and AD/mutant CRTC1^C216A/C216A^ hiPSCs, we transfected 6-week-old neurons with green fluorescent protein (GFP).

Approximately 5% of the cells expressed GFP, which allowed us to trace the length of neurites of the transfected cells. In line with our previous findings^41^, AD/CRTC1^WT/WT^ hiPSC neurons exhibited shorter total neuritic processes compared to isogenic non-AD/CRTC1^WT/WT^ hiPSC neurons **(Figure 4e).** However, we found that expression of non-nitrosylatable mutant CRTC1^C216A^ in the AD-hiPSC neurons significantly rescued this morphological defect (**Figure 4e**). This finding is consistent with the notion that SNO-CRTC1 contributes to the defect in neurite length in the AD-hiPSC neurons.

To assess apoptotic cell death, we performed terminal deoxynucleotidyl transferase (TdT) dUTP nick-end labeling (TUNEL) assays for DNA fragmentation in conjunction with condensed nuclear morphology on 6-week-old hiPSC neurons. We detected a significantly greater degree of cell death in AD/CRTC1^WT/WT^-hiPSC neurons compared to isogenic non-AD/CRTC1^WT/WT^-hiPSC neurons. Notably, cell death was totally abrogated in AD/CRTC1^C216A/C216A^-hiPSC neurons (**Figure 4f**).This experiment suggests that SNO-CRTC1 contributes to the cell death in the AD-hiPSC neurons.

### Non-nitrosylatable mutant CRTC1 rescues dysregulated gene expression in AD-hiPSC neurons

Next, we conducted several experiments to determine whether the therapeutic effect of non-nitrosylated CRTC1 was accompanied by an increase in CRTC1/CREB transcriptional activity. First, we performed a ChIP assay to examine whether non-nitrosylated CRTC1 retained the ability to bind to CREB in the promoter region of various downstream genes after KCl-induced depolarization. Anti-CRTC1 antibody (compared to negative control rabbit normal IgG antibody) was used to immunoprecipitate CRTC1/CREB-associated chromatin (**Extended Data Figure 6a**). We found significantly less recruitment of CRTC1 to the CREB-binding promoter region of the genes *Arc, cfos, EGR1*, and *BDNF* in AD/CRTC1^WT/WT^-hiPSC neurons compared to non-AD/CRTC1^WT/WT^-hiPSC neurons after KCl stimulation (**Figure 5a**). In contrast, non-nitrosylatable mutant AD/CRTC1^C216A/C216A^-hiPSC neurons demonstrated an increase in CRTC1 binding to the promoter region of these genes after KCl stimulation (**Figure 5a**).

**Figure 5.**
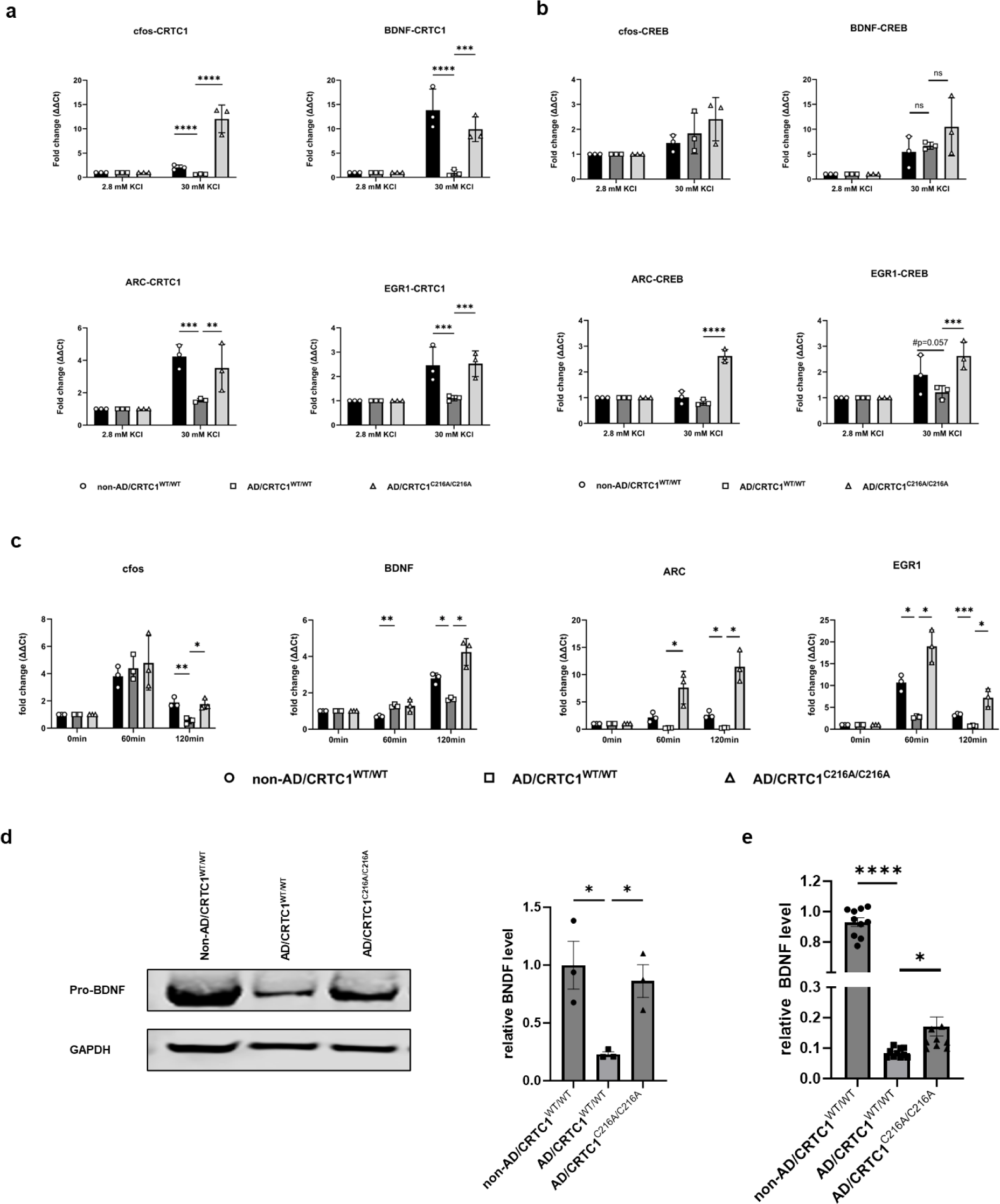
SNO-CRTC1 contributes to gene dysregulation and BDNF downregulation in hiPSC-derived AD models. **a,** Quantitative ChIP-qPCR analysis shows CRTC1^C216A/C216A^ mutation resulted in recovery of recruitment of CRTC1 to CRE in the promoter region of various downstream genes in hiPSC-derived AD neurons. hiPSC-derived neurons were treated with 30 mM KCl (or 2.8 mM KCl as a control) for 5 min, and then cells were collected and subjected to ChIP-qPCR for CRTC1 binding to the promoter region. Data represent mean fold enrichment (−ΔΔCT) ± SEM; n = 3 independent differentiations, analyzed by two-way ANOVA followed by Fisher’s LSD test, *p <= 0.05, **p < 0.01, ***p<0.001 and ****p < 0.0001. **b,** Quantitative ChIP-qPCR analysis of recruitment of CREB to the CRE region of downstream genes. hiPSC-derived neurons were treated and analyzed as in panel A except the ChIP-qPCR was for CREB binding to the promoter region. Data represent mean fold enrichment (−ΔΔCT) ± SEM; n = 3 independent differentiations, analyzed by two-way ANOVA followed by Fisher’s LSD test, *p < 0.05 and **P<0.01. **c,** Quantitative qPCR analysis shows that CRTC1^C216A/C216A^ mutation rescues CREB downstream gene transcription defects in hiPSC-derived AD neurons. Following a 5-min treatment with 30 mM KCl, mRNA was collected from hiPSC-derived neurons one and two hours later. Data represent mean fold enrichment (−ΔΔCT) ± SEM; n = 3 independent differentiations, analyzed by two-way ANOVA followed by Fisher’s LSD test, *p <= 0.05, **p < 0.01, ***p < 0.001, and ****p <0.0001. **d,** CRTC1^C216A/C216A^ mutation prevented downregulation of pro-BDNF protein in hiPSC-derived AD neurons. Cell lysates from 6-week-old hiPSC-derived neurons were analyzed by western blot. Data are mean ± SEM, and statistical differences were analyzed by two-way ANOVA and Tukey’s multiple comparisons post hoc test, n = 3 independent differentiations, *p < 0.05. **e,** CRTC1^C216A/C216A^ mutation prevented downregulation of mature BDNF protein in hiPSC-derived AD neurons. Conditioned medium from 6-week-old hiPSC-derived neurons was analyzed by ELISA with a specific antibody targeting the mature form of BDNF. Data are mean ± SEM, and statistical differences were analyzed by two-way ANOVA and Tukey’s multiple comparisons post hoc test, n = 3 independent differentiations, *p = 0.05 and ****p < 0.0001.

Prior studies had reported that recruitment of CREB to the promoter region of its target genes after neuronal stimulation was affected by NO^37^. To assess this effect in the context of AD neurons, we conducted a ChIP assay using a validated antibody against CREB (**Extended Data Figure 6b**). This assay allowed us to examine the recruitment of CREB and CRTC1 to the promoter region of interest. We found that CREB recruitment to the CRE element of c-*fos*, *Arc1,* and *BDNF* was not significantly different in our AD/CRTC1^WT/WT^-hiPSC neurons vs. our non-AD/CRTC1^WT/WT^-hiPSC neurons under basal conditions (**Figure 5b**). In contrast, after KCl stimulation, recruitment of CRTC1 to the same promoter region of these genes was significantly greater in AD/CRTC1^C216A/C216A^-hiPSC neurons than in non-AD/CRTC1^C216A/C216A^-hiPSC neurons or AD/CRTC1^WT/WT^-hiPSC neurons. Moreover, after KCl stimulation, CREB recruitment to the CRE element of the *ERG1* and *ARC* genes was similar to that of CRTC1 recruitment -- binding of CREB to the CRE element was decreased in AD/CRTC1^WT/WT^-hiPSC neurons compared to AD/CRTC1^WT/WT^-hiPSC neurons or AD/CRTC1^C216A/C216A^-hiPSC neurons (**Figure 5b**). In contrast, recruitment of CREB to the CRE element of the *cfos* and *BDNF* genes displayed a disparate pattern, with neuronal stimulation with KCl showing no inhibitory effect in non-AD/CRTC1^WT/WT^-hiPSC neurons compared to AD/CRTC1^WT/WT^-hiPSC neurons (if anything there was a trend toward increased CREB binding) (**Figure 5b**). In this case, AD/CRTC1^C216A/C216A^-hiPSC-neurons displayed either increased or no change in CREB binding (**Figure 5b**). This differential recruitment behavior of CREB among the various gene promoters may potentially be explained by the previous finding that CREB preoccupies the CRE element of some of its downstream genes prior to the initiation of neuronal activity^45^. Moreover, recent structural insights have revealed that CRTCs bind to both CREB and DNA^46,47^, and raise the possibility that disrupting CRTC binding via S-nitrosylation could also affect CREB-DNA binding in chromatin precipitation assays, at least for some genes.

Using quantitative qPCR, we next determined whether non-nitrosylatable CRTC1 could rescue transcription of genes downstream of CREB stimulation in AD/CRTC1^WT/WT^-hiPSC-neurons compared to AD/CRTC1^C216A/C216A^-mutant hiPSC-neurons. For this purpose, mRNA was collected from the hiPSC-derived neurons at baseline (0 min), or at 60 and 120 minutes after a 5-minute period of stimulation with 30 mM KCl, and monitored for transcription of *Arc, ERG1, BDNF,* and *cfos*. While the exact temporal pattern varied for each gene product, in general after neuronal stimulation, the mRNA for each of these CREB downstream genes was significantly greater in non-AD/CRTC1^WT/WT^-hiPSC neurons than in AD/CRTC1^WT/WT^-hiPSC neurons. Moreover, at any given time point, the mRNA values for the AD/CRTC1^C216A/C216A^-non-nitroyslatable mutant hiPSC-neurons showed significantly higher expression than the AD/CRTC1^WT/WT^-hiPSC neurons (**Figure 5c**).

### Non-nitrosylatable mutant CRTC1 prevents downregulation of BDNF in AD-hiPSC neurons

The CRTC1/CREB downstream gene, BDNF, has been shown to foster neurite outgrowth and neuronal cell survival^48–50^. However, BDNF is downregulated in human postmortem AD brains^51–53^. Similarly, when we monitored the intracellular level of pro-BDNF protein in 6-week-old hiPSC-derived neurons by immunoblot, we found significantly lower levels in AD/CRTC1^WT/WT^- than in isogenic non-AD/CRTC1^WT/WT^-hiPSC neurons. In contrast, BDNF levels in non-nitrosylatable mutant AD/CRTC1^C216A/C216^– hiPSC neurons were virtually completely rescued (**Figure 5d**).

We also measured the level of mature BDNF in conditioned medium (CM) from these hiPSC-derived neuronal cultures using an enzyme-linked immunosorbent assay ELISA).Three days after routine bi-weekly medium change, we found lower BDNF levels in AD/CRTC1^WT/WT^- than in AD/CRTC1^WT/WT^-hiPSC neurons, whereas the level of BDNF in the CM from the AD/CRTC1^C216A/C216A^ -hiPSC neurons was significantly increased compared to AD/CRTC1^WT/WT^-hiPSC neurons (**Figure 5e**).

### Non-nitrosylatable mutant CRTC1 reverses synapse loss in 5XFAD transgenic mice

CRTC1 has been reported to affect synaptic integrity, and the loss of synapses is arguably the best correlate to cognitive decline in human AD patients^54–56^. Therefore, to determine if SNO-CRTC1 contributes to synaptic loss *in vivo*, we used the 5XFAD transgenic mouse model of AD^33,57^. We injected lentiviral vectors expressing CRTC1 (WT CRTC1 or non-nitrosylatable mutant CRTC1^C216A^) or control vector that only expresses tdTomato into the hippocampus of 5XFAD mice (**Extended Data Figure 7a**). One month after injection, we assessed CRTC1 expression using tdTomato as a marker. In these experiments, there was no statistically significant difference in the expression level of WT and non-nitrosylatable mutant CRTC1 (**Extended Data Figure 7b**). Next, we examined histologically the effects of WT CRTC1 and non-nitrosylatable CRTC1^C216A^ on presynaptic integrity. Quantitative, unbiased confocal immunohistochemistry analysis of hippocampal sections stained with a monoclonal antibody against the presynaptic marker, synaptophysin (SY38), revealed that expression of CRTC1^C216A^, but not WT CRTC1, significantly increased the signal for SY38 compared to control littermate 5XFAD mice injected with control vector (**Figure 6a**). Although there was a trend toward improved presynaptic signal with WT CRTC1 overexpression, the value did not reach significance (**Figure 6b**); this result was similar to that found with other SNO-proteins affecting synaptic number, possibly because overexpression of WT protein can act as a sink for NO while leaving additional WT protein unnitrosylated^58,59^, and WT CRTC1 protein is known to be decreased in AD patient brain^30^.

**Figure 6.**
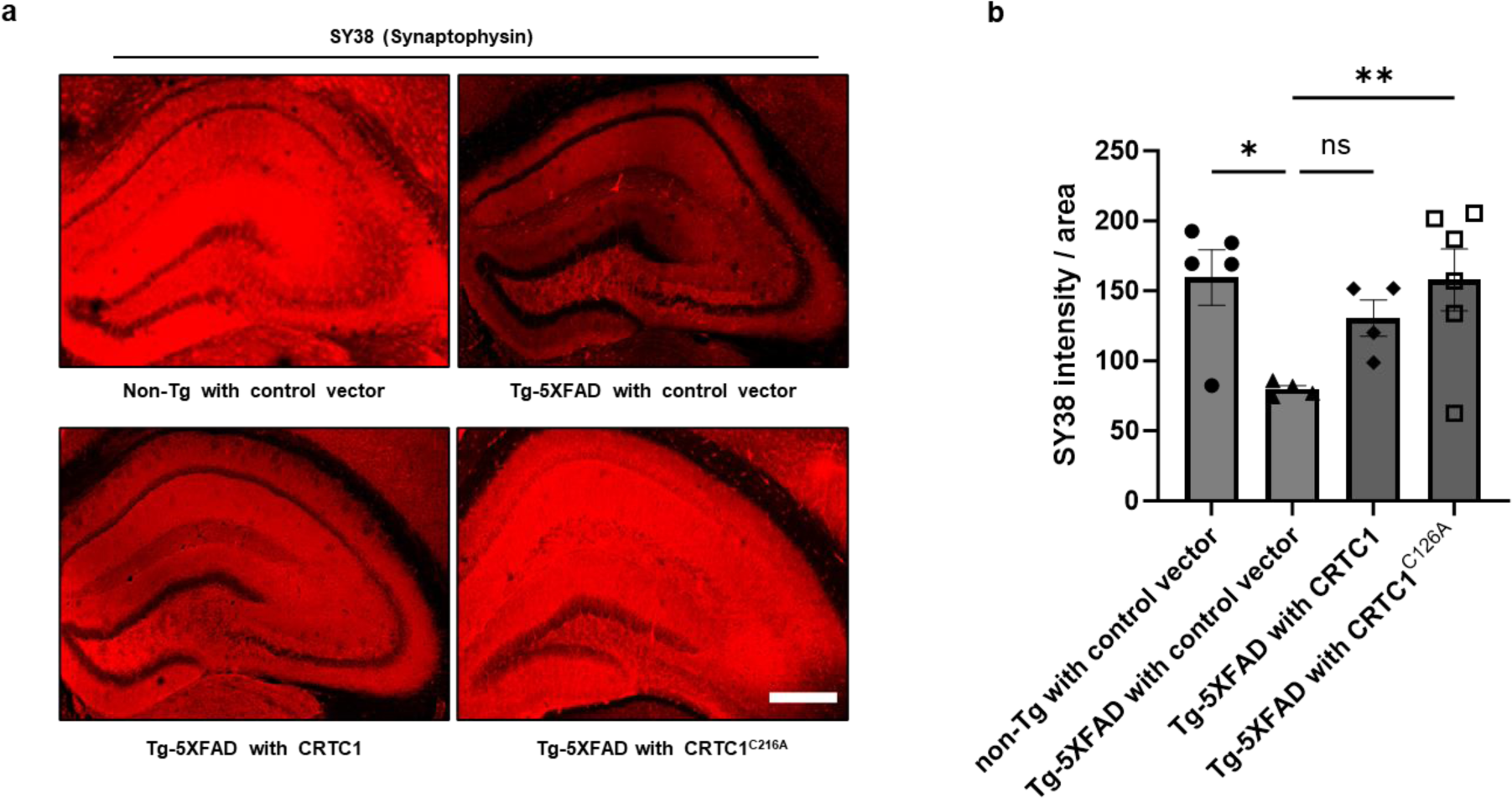
Lentiviral expression of CRTC1^C216A^ in vivo prevents loss of presynaptic marker in 5XFAD mice. **a,** Representative immunostaining of hippocampal sections with synaptophysin antibody (SY38). **b,** Quantification of hippocampal immunohistochemistry from WT and 5XFAD mice injected with WT, mutant CRTC1^C216A^ or control lentiviral vectors only express tdTomato. Immunohistochemistry was performed with synaptophysin antibody (SY38). Values are mean ± SEM; n = 5 for non-Tg/WT littermates injected with control vector; n = 4 for 5XFAD injected with control vector; n = 4 for 5XFAD injected with CRTC1; and n = 6 for 5XFAD injected with CRTC1^C216A^. *p < 0.05, **p < 0.01 by ANOVA with Fisher’s post hoc LSD test. Scale bar, 200 µm.

### Non-nitrosylatable mutant CRTC1 rescues LTP in 5XFAD mice

Studies have shown that 5XFAD mice exhibit reduced LTP magnitude and impaired synaptic transmission compared to WT mice^33^. It has previously been shown that CRTC1 is essential for the induction and maintenance of LTP and late phase LTP (L-LTP)^60–63^. Here, we investigated the effects of overexpressing non-nitrosylatable mutant CRTC1^C216A^ vs. control vector on LTP in 5XFAD mice aged 6-9 months. For this purpose, we injected lentiviral vectors expressing these plasmids into the hippocampus of 5-7-month-old 5XFAD mice. One month after injection, we performed electrophysiological assessments of synaptic plasticity on hippocampal slices isolated from the ipsilateral hippocampus. We evaluated the magnitude and the time course of LTP of synaptic transmission between CA3 and CA1 neurons, as described in the Methods. We assessed changes in field excitatory postsynaptic potentials (fEPSP) slope as an indicator of synaptic plasticity, and expressed LTP as relative values compared to baseline prior to stimulation and the potentiated fEPSP.

In our experiments, theta bursts reliably induced LTP of the fEPSP in all groups. The initial post-tetanic potentiation (PTP) was characterized by rapid decay that lasted <15 min and transitioned to stable LTP that lasted well over 1 hr. For comparisons between groups, we looked at the mean LTP magnitude obtained at 30-35 min. In WT littermate mice injected with control vector, LTP was 179.7 ± 15.7% of the baseline (n =11) and was comparable to the 5XFAD group injected with non-nitrosylatable mutant CRTC1^C216A^ (192.5 ± 8.7%, n = 13). However, the magnitude of LTP in slices from 5XFAD mice injected with control vector was significantly impaired at 122.8 ±5.6% of the baseline (n = 11, ANOVA, *F*_3,45_ = 9.057, p < 0.001).

Overall therefore, we observed that theta burst-induced LTP was significantly impaired in brain slices from 6 to 8-month-old Tg 5XFAD mice injected with a control vector when compared to age-matched WT (non-Tg) mice injected with the same vector. Notably, overexpression of non-nitrosylatable CRTC1 rescued LTP in the 5XFAD mice to the same level as that observed in WT mice (**Figures 7a and b)**. These findings provide compelling evidence supporting the notion that non-nitrosylatable CRTC1 plays a critical role in modulating LTP in the 5XFAD mouse model of AD. Collectively, these results are consistent with the notion that SNO-CRTC1 not only contributes to dysregulation of neuronal CREB-dependent gene expression, but also the ensuing contribution of these genes to neuronal form, function, and synaptic plasticity.

**Figure 7.**
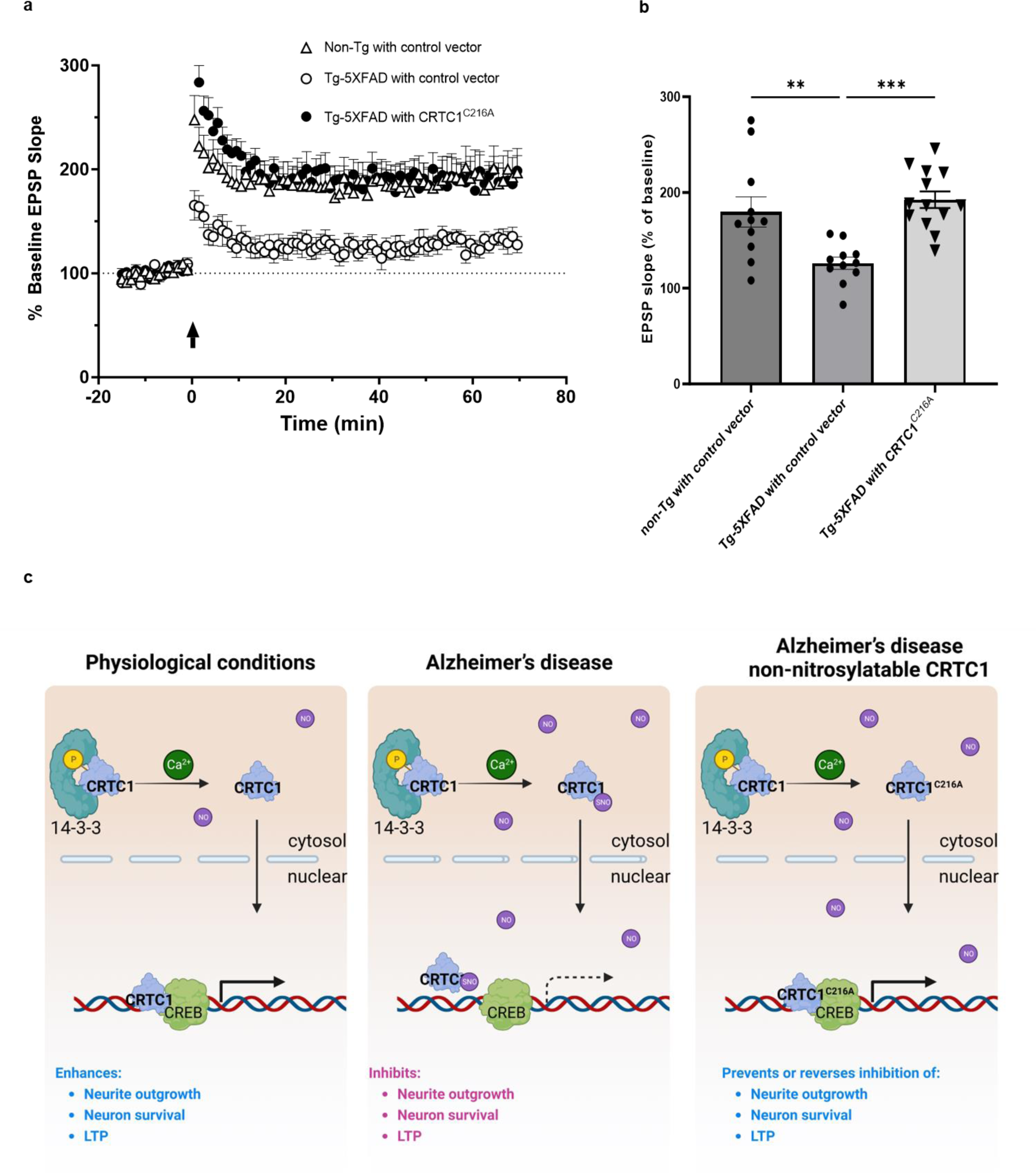
Non-nitrosylatable CRTC1 rescues LTP defects in 5XFAD mice. **a,** LTP data from non-nitrosylatable mutant CRTC1-injected AD transgenic mice. Non-nitrosylatable mutant CRTC1^C216A^ (or control vector) was delivered via lentiviral injection into the hippocampus of 5XFAD mice, followed by LTP assessment one month later. **b,** Group comparisons and statistical evaluations of LTP results performed at 30-35 minutes after LTP induction. Values are mean + SEM; n = 11 for non-Tg/WT littermates injected with control vector; n=11 for 5XFAD mice injected with control vector; and n = 13 for 5XFAD mice injected with CRTC1^C216A^. **p < 0.01 and ***p < 0.001 by ANOVA with Dunnett test. **c,** Schema for action of S-nitrosylated CRTC1 in CREB-mediated gene activation. Under normal conditions, CRTC1 translocates into the nucleus and interacts with CREB to stimulate gene transcription in response to neuronal activity. However, in neurodegenerative disorders such as AD, generation of excessive NO leads to the formation of SNO-CRTC1, disrupting the interaction of CRTC1 with CREB and thus impairing gene expression. Consequently, crucial processes in neurons are disrupted, including neurite outgrowth, synaptic plasticity (in the form of LTP), synapse maintenance, and neuronal cell survival. The introduction of a non-nitrosylatable CRTC1 mutant largely restores the normal gene expression pattern and mitigates against these defects in AD models both *in vitro* and *in vivo*.

## DISCUSSION

Disruption of CRTC1/CREB-dependent gene expression has been implicated in the pathogenesis of AD and is thought to be associated with Aβo accumulation^10,11,30,64^. Although the abundance of CRTC1 protein is reduced in AD patients’ brains with intermediate and advanced disease (Braak stages IV-VI), dysregulation of CRTC1-mediated gene transcription has been observed prior to that time, even at early stages of the disease (Braak stages I–II)^30^, as well as in early stages in AD transgenic mouse models^10,11,30^. Accordingly, additional mechanisms must be involved in dysregulation of CRTC1-mediated gene transcription in early AD. Along these lines, increased nitrosative stress due to generation of excessive levels of NO has been shown to occur with normal aging as well as in neurodegenerative diseases, and protein S-nitrosylation in the brain is elevated starting at early stages in both AD mouse models and AD human brains^9,65–67^. In this study, we provide evidence that increased levels of NO-related species result in S-nitrosylation of CRTC1 at Cys216 (forming SNO-CRTC1). Additionally, we demonstrate that the formation of SNO-CRTC1 interferes with the interaction of CRTC1 and CREB, thus compromising neuronal activity-dependent expression of downstream CRTC1/CREB-regulated genes.

Furthermore, we utilized CRISPR/Cas9 technology to knock-in non-nitrosylatable mutant CRTC1^C216A/C216A^ into an AD hiPSC line in which one allele bears the APPSwe mutation of familial AD. We found that activity-induced CRTC1/CREB gene transcription was restored in neurons derived from the non-nitrosylatable CRTC1 AD-hiPSC line compared to AD hiPSC-derived neurons bearing WT CRTC1. Additionally, non-nitrosylatable mutant AD/CRTC1^C216A/C216A^-hiPSC neurons displayed restoration of neurite outgrowth compared to the stunted dystrophic neurites of AD/CRTC1^WT/WT^-hiPSC neurons, and CRTC1^C216A^ protected AD hiPSC neurons from cell death. Moreover, we show that non-nitrosylatable mutant CRTC1 rescued dysregulated CREB-directed gene expression, including BDNF expression, which is known to influence neurite outgrowth and neuronal survival, in this AD hiPSC-derived neuronal model system.

Finally, we demonstrate that overexpression of non-nitrosylatable mutant CRTC1^C216A^ *in vivo* in the 5XFAD mouse model can prevent synapse loss, which is known to underlie cognitive decline in AD^54–56^. Additionally, *in vivo* expression of non-nitrosylatable mutant CRTC1^C216A^ prevented loss of LTP in 5XFAD mice, indicating that synaptic plasticity was preserved in the absence of SNO-CRTC1 formation and its consequent inhibition of CREB-mediated gene transcription. Taken together, based on both rodent and human *in vitro* and *in vivo* AD model systems, these multiple lines of evidence are consistent with the notion that formation of SNO-CRTC1 contributes to dysregulated CREB-mediated gene expression, with consequent neuronal and synaptic damage resulting in aberrant synaptic plasticity in the context of AD.

A potential caveat of the *in vivo* results might involve the cytoplasmic chaperone of CRTC1, 14-3-3 protein. It is possible that overexpression of CRTC1 could outpace the ability of 14-3-3 to act as a chaperone for CRTC1 and therefore alter its function^46^. It is unlikely that this was an issue here, however, because we observed similar levels of expression of mutant and WT CRTC1 in brains injected with these constructs, and only the non-nitrosylatable mutant manifested a significant neuroprotective effect on synaptic density. Collectively, our results demonstrate that SNO-CRTC1 may play a key role in AD pathogenesis.

Emerging evidence suggests that a large number of cytoplasmic and nuclear proteins are regulated by S-nitrosylation, with many of them involved in regulation of gene expression^36^. Previous work suggested that NO influences the binding of CREB to DNA and thus the expression of CRE-regulated genes in a phosphorylation-independent manner. S-Nitrosylation was reported to enhance CREB binding to CRE sites located in the promoter regions of *C-FOS* and *ERG1*, representing immediate early genes (IEGs)^36–38,68^. Many of the IEGs are transcription factors themselves and are believed to initiate secondary programs of gene expression that are involved in the regulation of neurite outgrowth, memory formation and neuron survival^69,70^. However, these prior studies of NO effects on IEG expression were not conducted in the context of a neurodegenerative disease model or with neuronal stimulation. In contrast, the present findings indicate that an excessive level of NO, as found in AD and in our AD-related models, including rat primary cerebrocortical neurons exposed to Aβo, AD patient hiPSC-derived cerebrocortical neurons, and 5XFAD and hAPP-J20 Tg mice, was associated with increased formation of SNO-CRTC1. We show in this disease context that S-nitrosylation of CRTC1 disrupted its binding to CREB, leading to dysregulated activity-dependent gene transcription. In our ChIP-assays using AD hiPSC-derived neurons, we found evidence that excessive levels of NO affect CRTC1 and CREB differentially under disease conditions. Specifically, pathological levels of NO did not significantly decrease CREB binding to the CRE elements of c-fos and Arc as well as BDNF, but did inhibit CRTC1 binding via S-nitrosylation. Additionally, in AD-related models, Aβo primarily trigger NO generation through enhanced activation of extrasynaptic NMDA receptors (eNMDARs)^40^. Calcium entry mediated by eNMDARs has been shown to activate a dominant CREB shut-off pathway, resulting in the blockade of BDNF expression induction^71^, which is also consistent with our findings for a role of SNO-CRTC1 in this effect.

Concerning cellular distribution, CRTC1 is an intrinsically disordered protein whose location and function is heavily dependent on PTMs and binding partners^72^. In resting neurons, CRTC1 is heavily phosphorylated, which facilitates its interaction with cytoplasmic 14-3-3 proteins at synapses. In response to neuronal activity, CRTC1 is dephosphorylated and translocated into the nucleus in a calcium- and cAMP-dependent manner^24^. We observed that acute exposure of rat cerebrocortical neurons to an NO donor dramatically increased the accumulation of CRTC1 in the nucleus. However, this effect was accompanied by a large increase in intracellular Ca^2+^, which may have triggered this nuclear translocation of CRTC1, as NO is known to transiently disrupt intracellular calcium homeostasis^35^. In fact, we observed that CRTC1 returned to the cytoplasm in rat primary cerebrocortical neurons an hour after exposure to the physiological NO donor SNOC, a time when SNO-CRTC1 levels still remained very high. This result suggests that S-nitrosylation of CRTC1 is not correlated with its distribution in the cell cytoplasm or nucleus and is consistent with our aforementioned results that S-nitrosylation primarily regulates CRTC1 binding to CREB/CRE.

Previous studies reported that CRTC1 plays a significant role in regulating BDNF gene expression^73–75^. A number of AD models have demonstrated the protective effects of BDNF on neuronal survival, synaptic density, and cognitive function^76–78^. Here, we found that SNO-CRTC1 inhibited activity-dependent BDNF expression under AD-related pathological conditions. In contrast, preventing SNO-CRTC1 formation with the NOS inhibitor L-NAME or using CRISPR/Cas9 to introduce non-nitrosylatable mutant CRTC1 significantly prevented downregulation of BDNF expression in these AD model systems. These findings suggest that SNO-CRTC1-mediated downregulation of BDNF expression may have contributed to the observed decrease in synaptic density that we observed in 5XFAD mouse and that was rescued by injection of non-nitrosylatable mutant CRTC1^C216A^. Activity-dependent transcription of other CRTC1-dependent CRE-containing genes associated with memory (*c-fos, Arc*, and *Erg1*) was also impaired in our AD-related model systems when excessive levels of SNO-CRTC1 were present and were reversed by reducing S-nitrosylation of CRTC1.

In summary, we show in various AD model systems, including hiPSC-derived AD neurons and AD transgenic mouse models, that formation of SNO-CRTC1 disrupted neuronal activity-dependent gene transcription mediated by CREB as well as neurite length and synaptic health. These results in the context of the finding that SNO-CRTC1 occurs at early stages of disease in transgenic AD mouse models lead us to speculate that S-nitrosylation of CRTC1 may contribute to early synaptic loss in AD brain. Since synaptic loss is arguably the best correlate of cognitive decline^54–56^, these findings suggest that SNO-CRTC1 represents a possible therapeutic target for early intervention in AD. Collectively, our results provide mechanistic insight for protein S-nitrosylation regulating activity-dependent gene expression in cerebrocortical neurons via formation of SNO-CRTC1, thus contributing to the pathogenesis of AD. Moreover, the findings have important therapeutic implications for AD, suggesting a novel pathway for disease modification.

## METHODS

### Culture of cell lines and media

Human neuroblastoma cells (SH-SY5Y; ATCC, CRL-2266) were cultured in DMEM (Thermo Fisher Scientific) supplemented with 10% fetal bovine serum (Gibco), 2 mM L-glutamine (Thermo Fisher Scientific), and 1% penicillin-streptomycin (Thermo Fisher Scientific). For depolarization of cells, a high-K^+^ solution was used (composition: KCl 30 mM, NaCl 112.8 mM, HEPES 10 mM, MgCl_2_ 1 mM, CaCl_2_ 1 mM and glucose 10 mM; pH 7.4) compared to physiological medium (containing KCl 2.8 mM, NaCl 140 mM, HEPES 10 mM, MgCl_2_ 1 mM, CaCl_2_ 1 mM, and glucose 10 mM, pH 7.4).

### Primary rat cerebrocortical cultures

Primary cerebrocortical cells were harvested from E16 Sprague-Dawley rats and maintained in culture as described previously^58,79^. Briefly, dissected cerebrocortices were digested with trypsin to make dissociated cells, which were then plated on poly-L-lysine-coated plates in D10C medium (80% high glucose DMEM (Thermo Fisher Scientific), 10% bovine calf serum, 10% F12 (HyClone), 30 mM HEPES (Omega), 2 mM L-glutamine (Gibco), and 0.25% penicillin-streptomycin. Experiments were performed 14-21 d after plating. For SNOC exposure, cerebrocortical cultures were exposed to100 μM of freshly prepared SNOC in DMEM medium without serum, and cells were collected 10 min later. To induce NO production by Aβo, cerebrocortical cultures were exposed to 500 nM Aβo_1-42_ in conditioned D10C medium for 3 hours, as we have described^39^.

### Differentiation of hiPSCs into cerebrocortical neurons

Differentiation of hiPSCs was performed as previously described with minor modifications^41^. Feeder-free hiPSCs were routinely maintained on Matrigel in mTeSR 1 medium (StemCell Technologies). Differentiation of hiPSCs to hNPCs was started with induction of neurospheres in aggrewell plates (StemCell Technologies) and exposure to a cocktail of small molecules containing 2 μM each of Dorsomorphin (Tocris), A83-01 (Tocris) and PNU74654 (Tocris) for 6 days in DMEM/F12 medium containing 20% Knock Out Serum Replacement (Invitrogen), 1x non-essential amino acids (Thermo Fisher Scientific) and 1x β-mercaptoethanol (Thermo Fisher Scientific). Neurospheres were transferred to Matrigel-coated plates and cultured in Neural Medium (DMEM/F12 medium supplemented with N2, N2, and 20 ng ml^-1^ of basic FGF (R&D Systems Inc.)) for two weeks to generate a monolayer of hNPCs. hNPCs were characterized by immunocytochemistry for hNPC markers PAX6, Nestin and SOX2. For terminal differentiation into cerebrocortical neurons, hNPCs were treated with 100 nM compound E (EMD Millipore) in Neural Medium for 48 hours and then maintained in Neural Medium supplemented with 20 ng/ml each of hBDNF and hGDNF for 3 weeks. Starting at week 4 of terminal differentiation, neurons were maintained in BrainPhys medium (StemCell Technologies). Successful generation of cerebrocortical neurons was confirmed by immunocytochemistry for FOXG1 and MAP2, and neurons were used for subsequent experiments after 6 weeks of differentiation unless otherwise noted in the text.

### Design and construction of Cas9–crRNA–sgRNA ribonucleoprotein

The Cas9–crRNA–tracrRNA ribonucleoprotein (RNP) contains a *Streptococcus pyogenes-* derived Cas9 nuclease (IDT) and an RNA duplex made of a CRISPR RNA (CRISPR-Cas9 tracrRNA–ATTO™ 550) (IDT) and a CRISPR guide RNA (sgRNA) that is partially complementary to the crRNA. The sgRNA was designed using the online CRISPR Design tool of Benchling (https://www.benchling.com/crispr) to target the PAM sequence nearest to the targeted mutation site (see **Table S1** for sgRNA sequence). The crRNA was fluorescently labeled, enabling us to generate single cell colonies by fluorescence-activated cell sorting in The Scripps Research Institute Flow Cytometry Core, La Jolla, California. Construction of Cas9– crRNA–sgRNA ribonucleoprotein was conducted according to the manufactory’s manual. Briefly, to form the crRNA:sgRNA duplexes, crRNA and sgRNA were mixed at a 1:1 ratio and heated at 95 °C for 5 min, followed by a cool down to room temperature. The RNA duplexes were then incubated with Cas9 enzyme at room temperature for 5min to assemble the RNP complex.

### RNP single cell colony generation and genotyping

Electroporation with freshly assembled RNP complex was performed on hiPSCs using the Human Stem Cell Nucleofector Kit 1 (Lonza) in conjunction with the Amaxa Nucleofector 2b system (Lonza) according to the manufacturer’s protocol. Twenty-four hours after electroporation, cells were separated by FACS and cells with ATTOtm550 fluorescence were sorted into a 96-well plate at a ratio of 1 cell per 2 wells to ensure the formation of single cell colonies. The single cell colonies were allowed to expand for 2 weeks, then dissociated and replated into a well of a 24-well plate for expansion.

Genomic DNA from individual colonies was isolated using QuickExtract DNA extraction solution (Lucigen). Then, 200 ng DNA and 1 μl of each primer (at 10 μM concentration) were used in a 25 μl PCR reaction utilizing Accuprim Pfx Supermix (Thermo Fisher). The resulting PCR products were directly subjected to Sanger sequencing to confirm the mutation. Genomic DNA from colonies with the desired mutation were subjected to PCR reaction with primers flanking the 3 most probable off-target sites, and the PCR products were assessed by Sanger sequencing. Only the colonies without detected off-target editing were used for subsequent experiments.

### Immunocytochemistry

Cells were fixed with 4% paraformaldehyde (PFA) for 10 minutes, washed once with PBS, and blocked with 5% BSA and 0.3% Triton X-100 in PBS for 1 hour. Cells were incubated with primary antibody against CRTC1 (1:1000, Cell Signaling), FOXG1 (1:250; Abcam), Nestin (1:200; Abcam), MAP2 (1:200, Invitrogen), or SOX2(1:200, Abcam) at 4 °C overnight, and the appropriate Alexa Fluor (488, 555, 647)-conjugated secondary antibody was used at 1:1000. For visualization of nuclear morphology, cells were counterstained with DAPI (1:500, Invitrogen). Images were collected by laser scanning confocal fluorescence microscopy (Nikon C2) by an observer masked to experimental group.

### Neurite tracing

hiPSC-derived neurons after 38-42 days of terminal differentiation were transfected with pCAG-MaxGFP (GFP) using Lipofectamine 2000 (Invitrogen). The GFP-labeled neurons were imaged at 10x magnification using a Nikon C2 confocal fluorescence microscope. Neurite tracing was performed using the neurite tracer plugin available in Image J. An observer masked to the experimental group traced the neurites for subsequent quantification.

### Biotin-switch assay

Biotin switch assays were performed as previously described^31,58^. Cells or tissues were lysed in blocking buffer containing 10 mM MMTS (S-methyl methanethiosulfonate, Sigma-Aldrich), 2.5% SDS, 100 mM HEPES (bioWORLD) at pH 7.4, 1 mM EDTA (Sigma-Aldrich) and 0.1 mM neocuproine (Sigma-Aldrich) and then incubated at 45 °C for 30 minutes to block free thiol groups. Next, 20 mM ascorbate (Sigma-Aldrich) was used to specifically reduce protein S-nitrosothiols and generate free thiols, which were subsequently biotinylated with 1 mM biotin-HPDP (Dojindo Laboratories). Finally, High Capacity NeutrAvidin agarose beads (Thermo Fisher Scientific) were used to pull down the biotinylated proteins, and the proteins were analyzed by immunoblotting using 5% of total protein for the loading control. The remaining protein was used for the NeutrAvidin agarose pull-down.

### Immunoblotting

Protein samples were separated on Bolt Bis-Tris Plus gels (Thermo Fisher Scientific) and analyzed on Immobilon-FL PVDF membranes (EMD Millipore). Membranes were first blocked with Odyssey TBS blocking buffer (Li-Cor), and then incubated with primary antibodies against CRTC1 (1:1000, Cell Signaling), CREB (1:1000, Cell Signaling), FLAG (1:5000, Sigma-Aldrich), GAPDH (1:5000, MilliporeSigma), or nNOS (1:1000, Santa Cruz). IR-dye 680LT-conjugated goat anti-mouse IgG (1:15000, Li-Cor) and IR-dye 800CW-conjugated goat anti-rabbit IgG (1:15000, Li-Cor) were used as secondary antibodies. The membranes were visualized and quantified using an Odyssey imaging system (Li-Cor).

### Detection of NO-related species by Griess assay

NO generation was monitored by Griess assay following the manufacturer’s instructions (Cayman) in combination with nitrate reduction, allowing us to measure the amount of total nitrite and nitrate, the major conversion products of NO in cells. An aliquot of tissue lysate was incubated with nitrate reductase and nitrate reductase cofactor for 1 hour at room temperature. Then the mixture was reacted with Griess reagent at room temperature for 20 minutes to generate a pink-red colored azo dye, which was quantified at 540 nm using an Optimax microplate reader (Molecular Devices).

### RT-qPCR

After various cell treatments, RNA was isolated using the RNeasy Mini Kit (QIAGEN). cDNA was synthesized using QuantiTect Reverse Transcription Kit (Qiagen), and RT-qPCR was performed using SYBR GREEN PCR master Mix (Thermo Fisher) on a QuantStudio 3 real-time PCR instrument (Applied Biosystems). For each reaction, there were 200 ng cDNA, 1 µl of each primer (10 µM concentration) and 1×SYBR mixture. Fold change was calculated using the comparative cycle threshold method. All samples were run in triplicate.

### Chromatin immunoprecipitation assays

Chromatin immunoprecipitation (ChIP) assays were conducted using the ChIP-IT EXPRESS assay kit protocol (Active Motif, Carlsbad, CA). After exposure to Aβo, cells were cross-linked with 1% formaldehyde at room temperature for 10 min, washed with PBS, quenched with glycine, and lysed, followed by centrifugation at 2400*g* to generate a nuclear pellet. The nuclear pellet was resuspended in enzymatic shearing buffer to shear DNA into 200–1000 bp fragments. The resulting suspension contained a protein and chromatin mixture. Subsequently, 10% of the mixture was saved for the “input” sample, and the rest was incubated with magnetic beads and anti-CRTC1 antibody (Cell Signaling) for 4 hours at 4 °C. Immunoprecipitated DNA was then eluted from the beads, and protein-DNA cross-linking was reversed. Input and ChIP DNA were analyzed using qPCR with primers within the promoter region of the target genes. qPCR was performed using SYBR green master mix on a QuantStudio 3 real-time PCR instrument (Applied Biosystems). Fold enrichment (n-fold) was calculated using the comparative cycle threshold method. All samples were run in triplicate.

### Terminal deoxynucleotidyl transferase dUTP nick end labeling (TUNEL) assays

Terminal deoxynucleotidyl transferase dUTP nick end labeling (TUNEL) assay was used to assess apoptosis in hiPSC-derived cerebrocortical neurons after 6 weeks in culture. TUNEL assay was performed using the In-Situ Cell Death Detection Kit (TMR Red) (Roche) according to the manufacturer’s protocol. Briefly, cells were fixed with 4% PFA for 10 min at room temperature, washed once with PBS, permeabilized with ice cold PBS plus 0.1% Triton X-100 and 0.1% sodium citrate for 2 min on ice and washed 3 times with PBS. Cells were then incubated with TUNEL reaction mixture at 37 °C in an incubator for 1 hour. After TUNEL staining, cells were blocked and incubated with primary antibody against the neuronal marker MAP2 (1:200, Invitrogen) overnight at 4 °C. For visualization of nuclei, cells were counterstained with DAPI (1:500, Invitrogen). At least 5 random fields and a minimum of 500 cells were counted per coverslip by an observer masked to the experimental group.

### Production of lentiviral constructs, stereotaxic injection in the brain, and quantitative immunohistochemistry

CRTC1 (WT or non-nitrosylatable mutant CRTC1^C216A^) was subcloned into the lentiviral pCDH-EF1-dTomato-T2A-Kpn21 vector (System Biosciences) and used to generate active lentiviral particles for injection into animals^59^. Both tdTomato and CRTC1 were inserted downstream of the EF1 promoter in the expression construct and connected via a T2A self-cleaving peptide. The lentiviruses were packaged by the Genetic Perturbation Screening Core at the UF Scripps Institute in Jupiter, Florida. 5XFAD transgenic mice (#034848-JAX) or non-transgenic littermate control mice at 5-7 months of age were stereotactically injected with lentiviral vectors into the right hippocampus at a rate of 0.1 μl per min, with a total of 1 μl of lentivirus infused (titer=1e9 for each vector). The injection coordinates used relative to the Bregma were as follows: Anteroposterior (AP) -2.0 mm, mediolateral (ML) 1.7 mm, and dorsoventral (DV) 1.65 mm. The needle was left in place for 10 minutes after the injection was completed before retraction to ensure sufficient tissue penetration and prevent unwanted lentiviral contamination to other brain areas. Expression of CRTC1 constructs were confirmed by using RFP antibody against tdTomato (Rockland Immunochemical).

One to two months after lentiviral injection, mice were anesthetized and transcardially perfused with PBS, followed by 4% PFA. The brains were then removed and further fixed in 4% PFA for 48 hours at 4 °C before being stored in 1% PFA until analysis. Quantitative immunohistochemistry was performed as described^39^ using a mouse monoclonal antibody against Synaptophysin (Clone SY38, Millipore). Analysis was carried out on the entire hippocampal image using Fiji (ImageJ) software. All procedures described here were approved by the Institutional Animal Care and Use Committees at the institutions where the work was conducted.

### Electrophysiological assessment of LTP

One month after lentiviral construct injection, we performed *ex vivo* electrophysiological assessments of synaptic plasticity on hippocampal slices (400 µm thick) isolated from the hippocampus ipsilateral to the injection. Briefly, acute hippocampal slices were prepared in oxygenated, ice-cold sucrose-substituted artificial cerebrospinal fluid (aCSF) (composition in mM): 190 sucrose, 25 NaHCO_3_, 10 NaCl, 3 KCl, 1.25 NaH_2_PO_4_, 0.1 CaCl_2_, 1 MgSO_4_, 25 D-glucose, aerated with 95% O_2_ and 5% CO_2_ and adjusted to pH 7.4. Slices were cut using a vibrating-blade microtome (Leica VT1200S). After sectioning, slices were rinsed and stored for at least 2 hours in recording aCSF (composition in mM): 125 NaCl, 3 KCl, 25 NaHCO_3_, 1.25 NaH_2_PO_4_, 10 Glucose, 1 MgCl_2_, and 2 CaCl_2_, aerated with 95% O_2_ and 5% CO_2_. pH was adjusted to 7.4 with NaOH and osmolality to 300 mOsm with sucrose. Slices were transferred to a recording chamber on an upright microscope and continuously superfused with oxygenated recording aCSF for the duration of the experiment. We evaluated the magnitude and time course of LTP of synaptic transmission between CA3 and CA1 neurons, as described previously with minor modifications^80,81^. To evoke synaptic transmission, we used bipolar concentric electrodes (Rhodes Medical Instruments) to stimulate the Schaffer collaterals/commissural pathway with constant square current pulses (0.05 ms). fEPSPs were recorded with glass microelectrodes filled with saline (resistance ∼1 MΩ) and placed in the stratum radiatum of CA1 neurons. The stimulation intensity (stimulation current pulse amplitude) was adjusted to evoke approximately 50% of the fEPSP amplitude before the fEPSP became contaminated with a population spike (which is known to be associated with firing of CA1 neurons). Higher stimulation intensities were avoided. To determine the time course of LTP, the synaptic response was evoked and monitored at a stimulation rate of 0.033 Hz, and two consecutive fEPSPs were averaged. Thus, one averaged fEPSP per minute was saved and used for offline analysis. During recordings, a low-pass filter was set at 3 kHz. Additionally, all voltage traces were further digitally filtered "offline" at 1 kHz using the Clampfit built-in algorithm. After approximately 15 minutes of baseline recordings, LTP was triggered by theta stimulation consisting of 4 trains of 5 pulses at 100 Hz, separated by 200 ms, each delivered at the same stimulation intensity. LTP of the fEPSP was then recorded for at least another 65 minutes. As a measure of synaptic transmission, we evaluated changes in the fEPSP slopes (initial 10-80% negative linear segment of the fEPSP slope) and maximal negative fEPSP amplitudes. LTP was expressed as relative values (ratios) between the average 15-minute baseline value (100%) and the potentiated fEPSP. To compare differences between groups, LTP magnitudes were averaged at 30-35 minutes after LTP induction. These 5-minute LTP segments reflected the maintenance phase of LTP. Differences between groups were statistically evaluated by ANOVA.

### Aβo peptide preparation

Aβo were prepared as previously described^39^. Briefly, Aβ_1-42_ was suspended in hexafluoro-2-propanol and incubated for 2 h at room temperature to generate a 1 mM suspension. The solvent was evaporated in a hood overnight, and the precipitate was resuspended in DMSO to a concentration of 5 mM, which was stored at −80 °C until use. For oligomerization, the stock solution was diluted 10-fold in MEM (GIBCO) and kept at 4 °C for at least 24 hours before use. Extent of oligomerization was determined by dynamic light scattering^39^.

## QUANTIFICATION AND STATISTICAL ANALYSIS

To ensure rigor and reproducibility, all data presented in the figures were obtained from multiple biological replicates from separate samples or differentiations (n ≥ 3), and the number of replicates is indicated in the figure legends. The number of animals or cultures used in each experiment was determined from power analysis on prior data using similar techniques. Data are expressed as mean ± SEM. Statistical analyses were performed using GraphPad Prism software (v.9). Differences between multiple groups were evaluated using an ANOVA followed by an appropriate post-hoc test. Differences between two groups were evaluated using an unpaired Student’s t test. A p-value < 0.05 was considered to be statistically significant.

## ACKNOWLEDGMENTS

We would like to express our gratitude to Professor Marc Montminy (Salk Institute for Biological Studies) for his helpful discussions and for providing us with the CRTCs plasmids. We also extend our thanks to Dr. Robert M. Witwicki at the Genetic Perturbation Screening Core, UF Scripps Research, Jupiter, FL, for lentivirus packaging. This study was supported in part by California Institute for Regenerative Medicine (CIRM) training grant EDUC4-12811 (to X.Z.), National Institutes of Health (NIH) grants R01 AG061845, R61 NS122098 and RF1 NS123298 (to T.N.), AA013498, AA006420 and AA027700 (to M.R.), and R35 AG071734, RF1 AG057409, R56 AG065372, R01 AG078756, R01 AG056259, R01 DA048882 and DP1 DA041722 (to S.A.L.).

