## Extended Data for "S-Nitrosylation of CRTC1 in Alzheimer’s disease impairs CREB-dependent gene expression induced by neuronal activity"

**Extended Data** **Figures and Figure Legends**

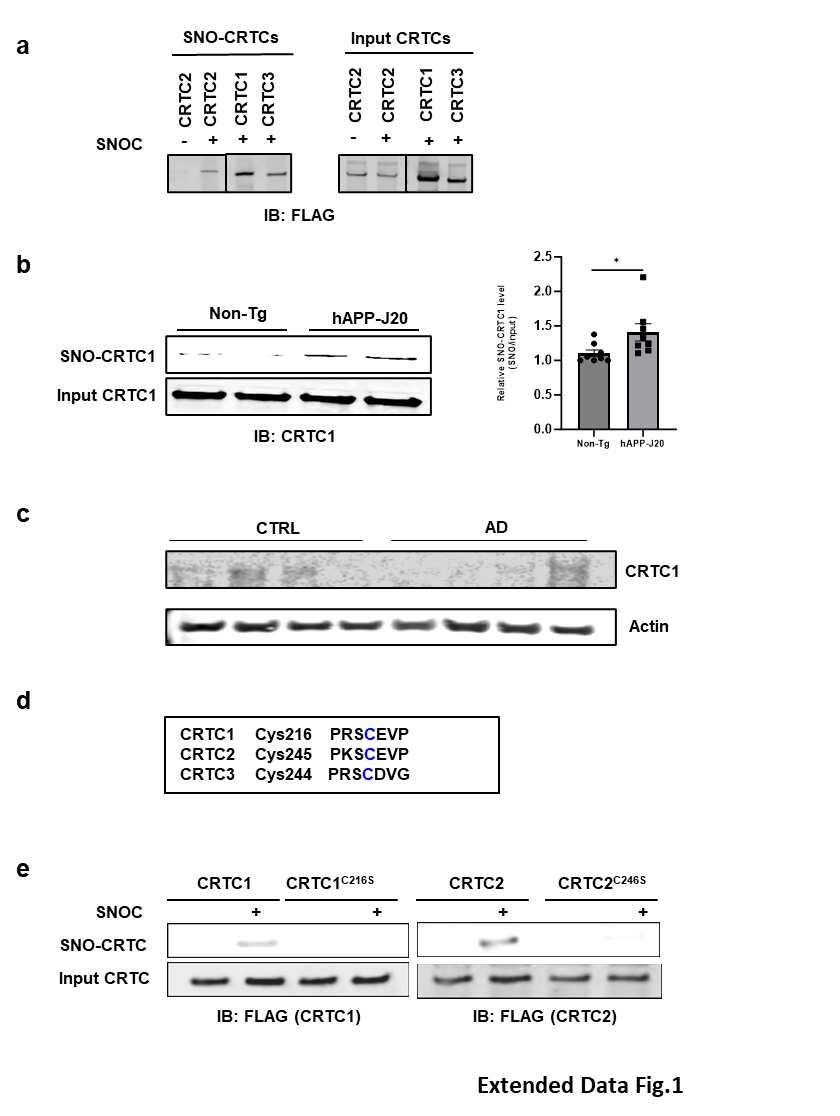

**Extended Data Figure 1 | All three isoforms of CRTCs are S-nitrosylated at a conserved cysteine residue**

**a,** HEK293 cells were transfected with FLAG-CRTC1-3 isoforms. After 1 day, cells were exposed to 200 μM fresh or old SNOC. Cell lysates were collected 10 min later, subjected to biotin-switch assay, and immunoblotted with anti-CRTC antibody.

**b,** Presence of SNO-CRTC1 in the cerebrocortex of J20-hAPP Tg AD mouse brains. Brain lysates from 3-month-old J20-hAPP or non-Tg/WT littermate mice were subjected to biotin-switch assay. Ratio of SNO-CRTC1/input-CRTC1 expressed as mean + SEM; n = 8 mice per genotype analyzed by two-tailed Student's t test, *p < 0.05.

**c,** Lack of substantial levels of CRTC1 protein in late-stage AD human postmortem brain or in age-matched controls.

**d,** Cys126 in CRTC1, Cys245 in CRTC2, and Cys244 in CRTC3 are the only conserved cysteine residues in the various CRTC isoforms.

**e,** The conserved cysteine residue is the major target of S-nitrosylation in CRTCs. HEK293 cells were transfected with FLAG-WT-CRTC1, FLAG-WT-CRTC2, mutant FLAG-CRTC1^C216S^, or mutant FLAG-CRTC2^C246S^. One day after transfection, cells were incubated with 100 μM fresh or old SNOC. Ten minutes later, cell lysates were subjected to biotin-switch assay and immunoblotted with anti-FLAG antibody. In this case, serine was substituted for cysteine in the non-nitrosylatable mutant and yielded similar results to the cysteine to alanine mutation used for other experiments (in general, alanine was used to avoid introducing a possible phosphorylation site in the mutant construct).

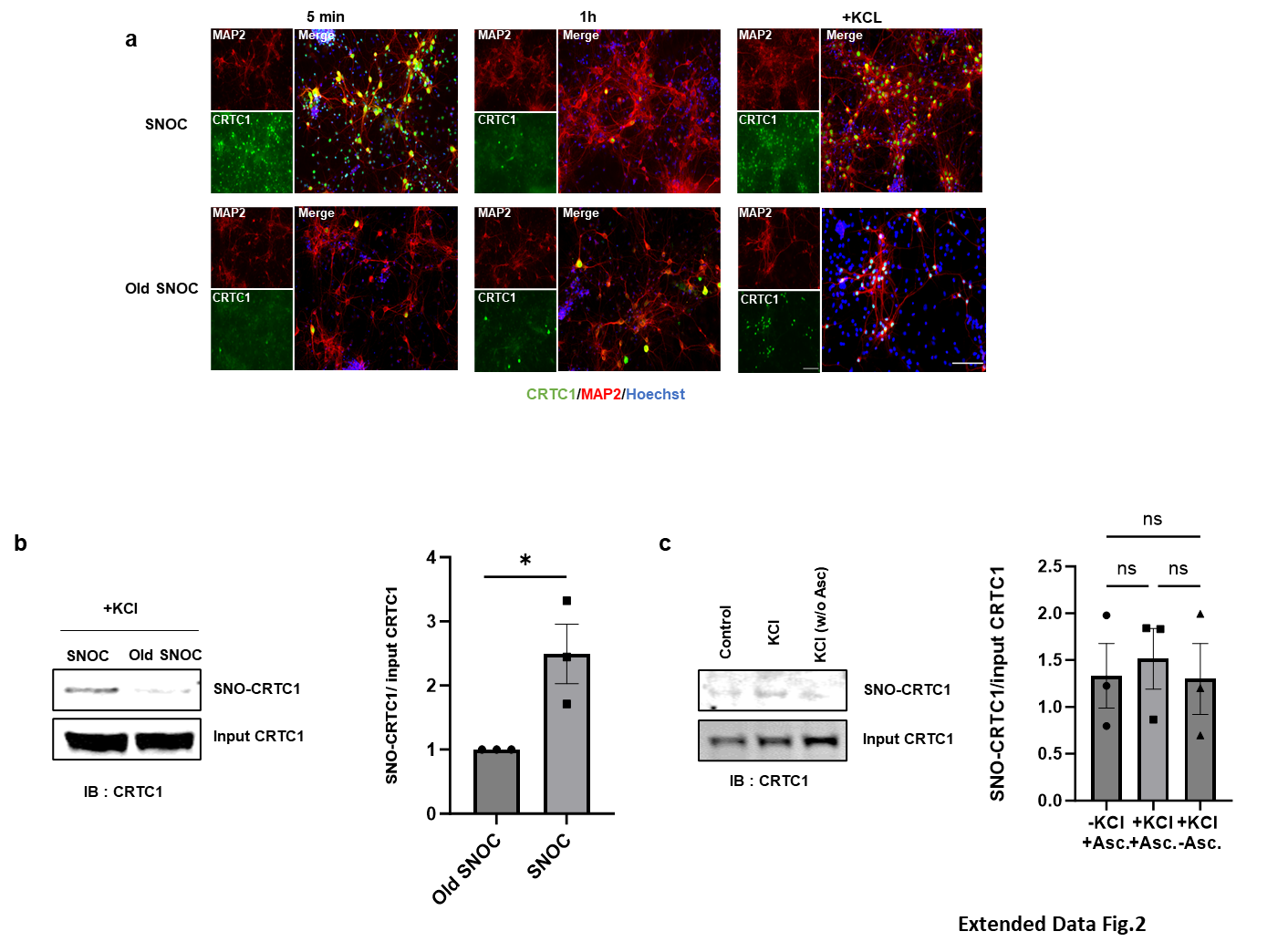

**Extended Data Figure 2 | S-Nitrosylation of CRTC1 does not affect KCl-induced nuclear translocation of CRTC1**

**a,** CRTC1 returned to the cytosol 1-hour after SNOC exposure. Primary rat cerebrocortical cultures were exposed to SNOC (100 µM); after 5 min, the cells were washed and put back into conditioned medium. One hour later, neurons were stimulated with KCl (30 mM) for 5 min. Cellular distribution of CRTC1 was detected by immunostaining: MAP2 (red); CRTC1 (green), and Hoechst nuclear dye (blue). Scale bar, 100 µm.

**b,** Formation of SNO-CRTC1 in primary rat cerebrocortical neurons 1 h after SNOC exposure. After SNOC or control (‘old‘ SNOC) exposure, cells were collected and subjected to biotin-switch assay. Differences between ratio of SNO-CRTC1/input CRTC1 were analyzed by two-tailed Student’s t-test, n = 3 independent experiments, *p < 0.05.

**c,** Depolarizing concentration of KCl (30 mM) does not induce SNO-CRTC1 formation. Primary rat cerebrocortical cultures were exposed to 30 mM KCl for 5 min, after which cells were collected and subjected to biotin-switch assay. SNOC-exposed samples in the absence of ascorbate (w/o Asc) served as a negative control. Values are mean ± SEM, n = 3 independent experiments, one-way ANOVA with Tukey’s multiple comparisons post hoc test, ns: not significant.

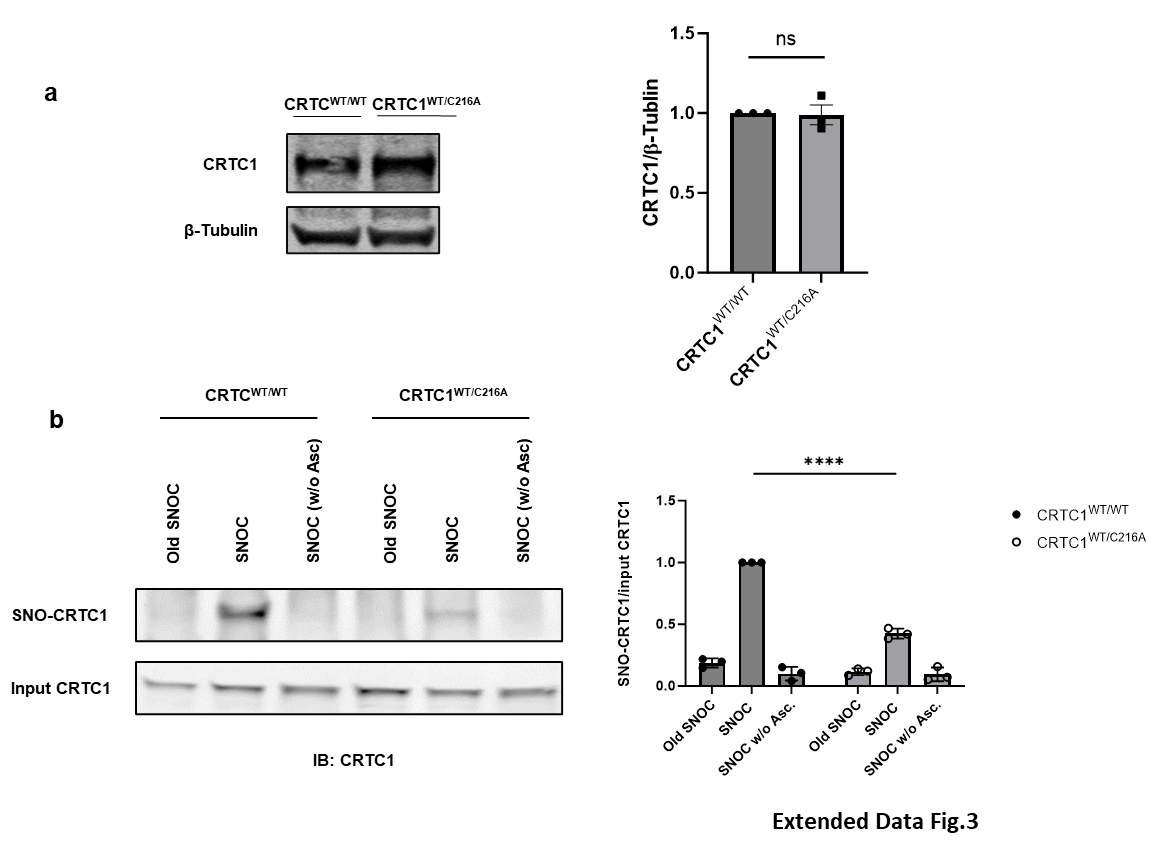

**Extended Data Figure 3 | Characterization of CRISPR/Cas9-engineered SHSY5Y cells heterozygous for non-nitrosylatable CRTC1 (SHSY5Y/CRTC1^WT/C216A^)**

**a,** CRTC1 levels were not changed in SHSY5Y/ CRTC1^WT/C216A^ by immunoblot. WT and mutant CRTC1^WT/C216A^ cells were collected, and cell lysates subjected to western blotting. Protein levels of CRTC1 were detected with anti-CRTC1antibody and normalized to protein level of β-tubulin. Differences between mean normalized protein levels of CRTC1 in WT and mutant cells (± SEM) were analyzed by two-tailed Student's t-test, n = 3 independent differentiations, ns: not significant.

**b,** Heterozygous non-nitrosylatable mutant CRTC1^WT/C216A^ significantly decreased SNO-CRTC1 formation relative to WT-CRTC1 IN SHSY5Y cells. Cells were collected and subjected to biotin-switch assay after exposure to SNOC. Values are mean ± SEM; n = 3 independent experiments, analyzed by two-way ANOVA followed by Tukey’s multiple comparisons post hoc test, ****p < 0.0001.

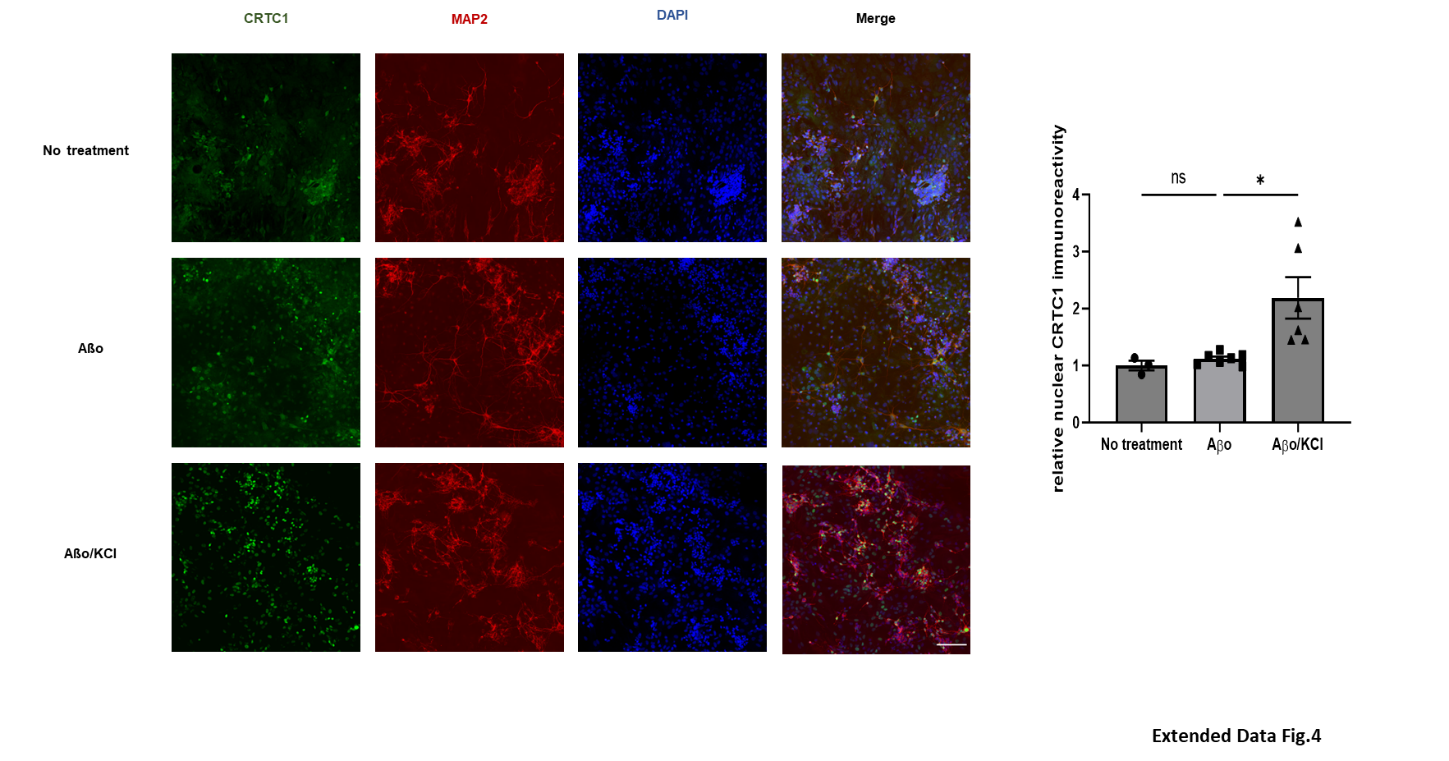

**Extended Data Figure 4 | Unlike incubation in conjunction with KCl, exposure to Aβo alone does not induce CRTC1 nuclear translocation**

Primary rat cerebrocortical cultures were exposed to 500 nM Aβo for 3 hours. Neurons were then fixed and immunostained with primary antibodies against MAP2 (red), CRTC1 (green), and Hoechst nuclear dye (blue). Quantitative CRTC1 nuclear immunoreactivity was analyzed by ANOVA followed by Tukey’s multiple comparisons post hoc test, *p < 0.05, ns: not significant. Scale bar, 100 µm.

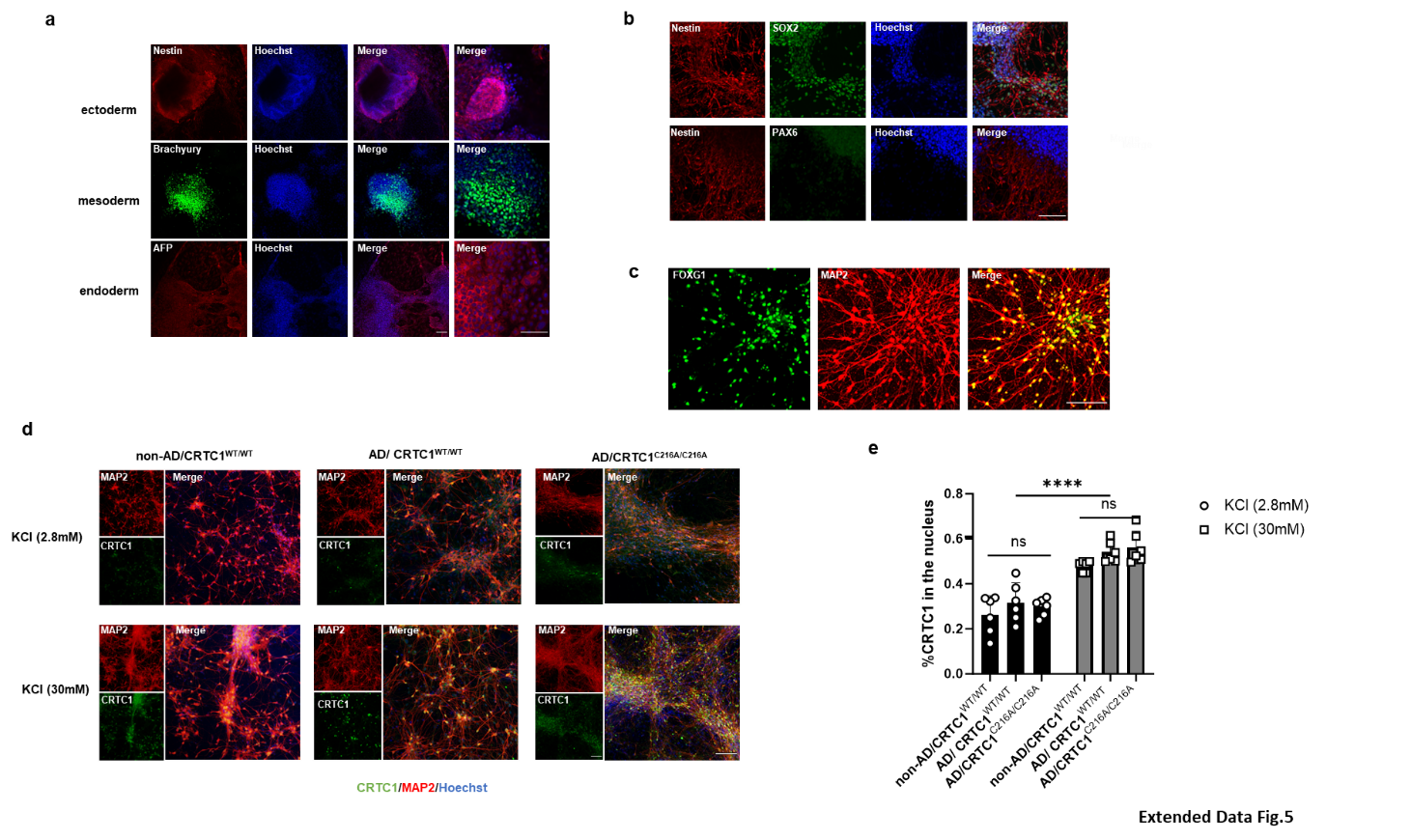

**Extended Data Figure 5 | Characterization of AD hiPSCs with non-nitrosylatable CRTC1 mutation introduced by CRISPR/Cas9**

**a,** Pluripotency of newly generated AD-CRTC1^C216A/C216A^ hiPSCs was investigated by embryoid body production using immunohistochemistry staining for specific germ-layer makers: Nestin for neuroectoderm, Brachyury for mesoderm and AFP for endoderm. Scale bars, 100 µm.

**b,** hNPCs generated from AD-CRTC1^C216A/C216A^ hiPSCs were characterized with specific markers. Top Panel: Nestin, PAX6, and Hoechst; bottom Panel: Nestin, SOX2, and Hoechst. Scale bar, 100 µm.

**c,** Characterization of cerebrocortical neurons derived from AD-CRTC1^C216A/C216A^ hiPSCs with markers FOXG1 and MAP2. Scale bar, 100 µm.

**d,** Non-nitrosylatable mutant CRTC1 does not affect KCl-induced nuclear translocation of CRTC1 in hiPSC-derived neurons. Cellular distribution of CRTC1 in hiPSC-derived neurons was examined by immunohistochemical staining after exposure to 30 mM KCl for 5 min using the following markers: MAP2 (red), CRTC1 (green), and Hoechst nuclear dye (blue). Scale bar, 100 µm.

**e,** Quantitative CRTC1 nuclear immunoreactivity was analyzed by ANOVA followed by Tukey’s multiple comparisons post hoc test, ns: not significant, , ****p < 0.0001.

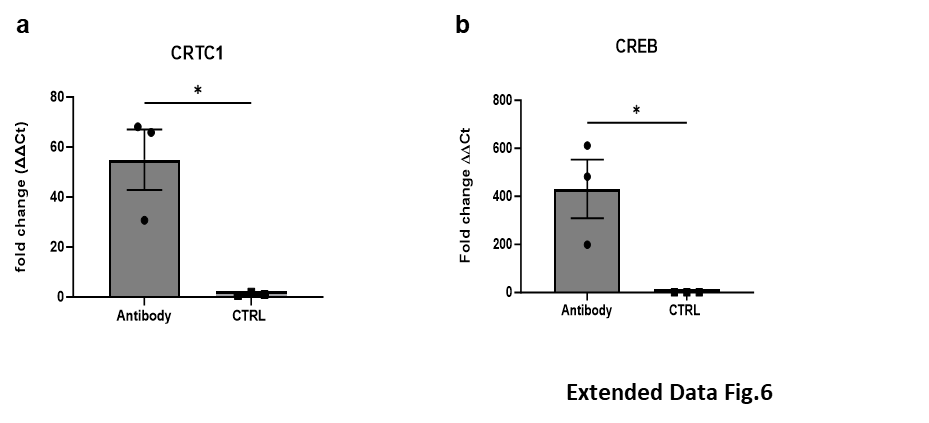

**Extended Data Figure 6 | Verification of specificity of anti-CRTC1 and anti-CREB antibodies used for CHIP-assay, related to Figure 5.**

**a,** Sheared chromatin was incubated with anti-CRTC1 antibody or negative control antibody (rabbit normal IgG) overnight at 4º C. Primers targeting the CRE promoter region of *c-fos* were used for CHIP-qPCR. Data represent mean fold-enrichment (−ΔΔCT) ± SEM; n = 3 technical replicates, analyzed by two-tailed Student’s t-test, *p < 0.05.

**b,** Sheared chromatin was incubated with anti-CREB antibody or negative control antibody (rabbit normal IgG) overnight at 4º C. Primers targeting the CRE promoter region of *c-fos* were used for CHIP-qPCR. Data represent mean fold-enrichment (−ΔΔCT) ± SEM; n = 3 technical replicates, analyzed by two-tailed Student’s t-test, *p < 0.05.

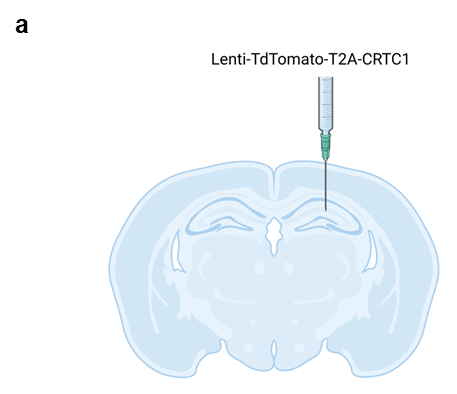

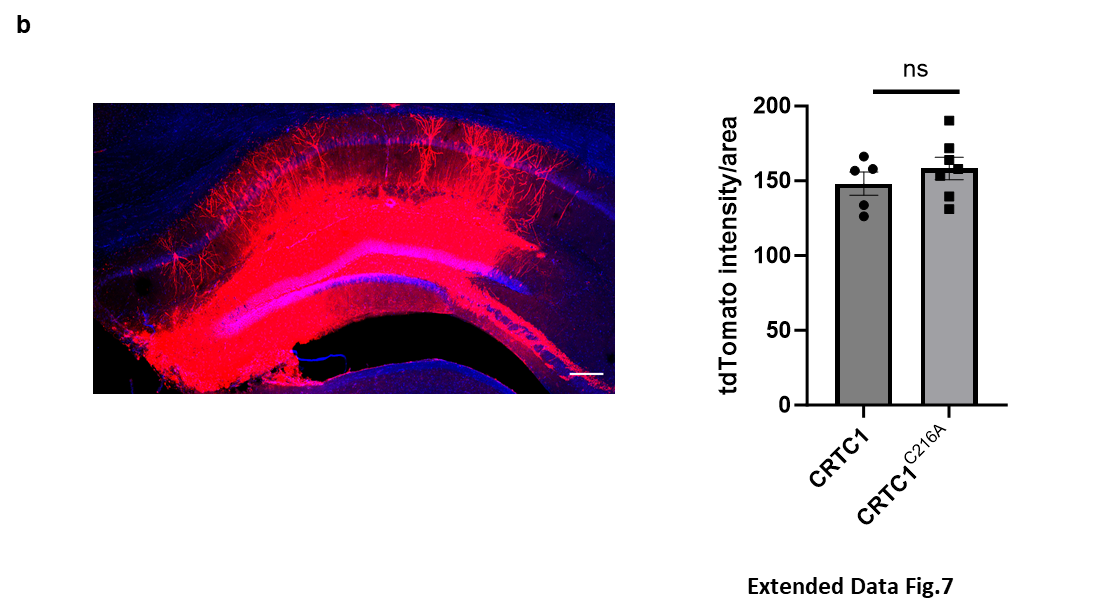

**Extended Data Figure 7 | Assessing expression of CRTC1 using tdTomato marker**

**a,** Diagram showing stereotaxic injection site of lentiviral constructs into the dentate gyrus. Both tdTomato and CRTC1 were inserted downstream of the EF1 promoter and connected via a T2A self-cleaving peptide.

**b,** One month after injection, we assessed expression of CRTC1 using tdTomato. Left: Representative image of tdTomato expression in the hippocampus of 5XFAD mice. Scale bar, 200 µm. Right: Data represent the mean fluorescence intensity over the entire hippocampus ± SEM; n = 4 for WT CRTC1, n = 7 for mutant CRTC1, analyzed by two-tailed Student's t-test, ns: not significant.
